# Engineering large-scale perfused tissues via synthetic 3D soft microfluidics

**DOI:** 10.1101/2021.08.20.457148

**Authors:** Sergei Grebenyuk, Abdel Rahman Abdel Fattah, Gregorius Rustandi, Manoj Kumar, Burak Toprakhisar, Idris Salmon, Catherine Verfaillie, Adrian Ranga

## Abstract

The vascularization of engineered tissues and organoids has remained a major unresolved challenge in regenerative medicine. While multiple approaches have been developed to vascularize in vitro tissues, it has thus far not been possible to generate sufficiently dense networks of small-scale vessels to perfuse large de novo tissues. Here, we achieve the perfusion of multi-mm^3^ tissue constructs by generating networks of synthetic capillary-scale 3D vessels. Our 3D soft microfluidic strategy is uniquely enabled by a 3D-printable 2-photon-polymerizable hydrogel formulation, which allows for precise microvessel printing at scales below the diffusion limit of living tissues. We demonstrate that these large-scale engineered tissues are viable, proliferative and exhibit complex morphogenesis during long-term in-vitro culture, while avoiding hypoxia and necrosis. We show by scRNAseq and immunohistochemistry that neural differentiation is significantly accelerated in perfused neural constructs. Additionally, we illustrate the versatility of this platform by demonstrating long-term perfusion of developing liver tissue. This fully synthetic vascularization platform opens the door to the generation of human tissue models at unprecedented scale and complexity.

## Introduction

Human engineered tissue and organoids are potentially transformational model systems, which could create dramatic efficiencies in the drug discovery process and function as key building blocks for regenerative medicine applications. In particular, larger-scale tissues have the possibility to recapitulate complex functional and organizational characteristics of their in vivo counterparts, and could therefore become a long-sought alternative to animal models^1–3^. However, the poorly defined structural organization, small size and slow maturation of these tissues have remained major limitations in engineering fully functional and reproducible organoids and tissues.

*In vivo*, the development of tissues is supported by a complex network of blood vessels which provide oxygen, nutrients and waste exchange and mediate paracrine interactions via growth and differentiation factors^4^. The size of the microvasculature is a critical parameter for local tissue perfusion: to maintain sufficient diffusion of oxygen, nutrients, and waste products most cells in vivo lie within 200 μm of a capillary. In the absence of vascular support, normal physiological conditions can be maintained only within this narrow range. Similar to the diffusion limits in normal tissue, the generation of solid tissue in vitro requires both vascularization and flow to maintain cell viability throughout the entire construct^1^. The lack of vascularization in engineered tissues therefore prevents oxygen and nutrient exchange, which is thought to be the main reason for the commonly observed development of a necrotic core within organoids once they reach a critical size, as well as for apoptosis within engineered tissues. These issues have been widely recognized, and various approaches have been reported in order to overcome them^5^.

The extrinsic induction of angiogenesis has been frequently used in the context of organoid vascularization. Vessel sprouts have been shown to infiltrate organoids maintained with endothelial cells in separate compartments of microfluidic culture devices^6, 7^, or co-cultured with pre-established microvascular beds^8, 9^. These results have suggested that the presence of a perfusable vasculature can enhance organoid growth^8^, confirming the importance of systemic cross-talk between vasculature and organoids in the developmental process. The resulting organoids have nonetheless been limited in size, and to date, functional organoid vascularization has only been achieved by grafting organoids into host animals^10, 11^.

A number of studies have focused on creating artificial vessels though the use of templating approaches based on patterned layer-by-layer deposition of gelled material, in the form of thin filaments^12^, droplets^13–18^ or layer-by-layer polymerization by stereolithography^19–21^. Large tissue constructs have been generated by bioprinting of bioinks comprised of gels carrying different cell types^12, 22, 23^, with vascular templates generated by depositing endothelial cells interleaved with tissue-specific cells using filament extrusion^24–28^ or stereolithography^19, 21^. While multiphoton lithography has been used to fabricate capillary-sized tubular fragments^29^ and vascular mimics^30^, these vessels have thus far not been successfully perfused. The dissolution of sacrificial networks to form lumenized vessels has been proposed as an alternative strategy, with resorbable gel filaments being created via stereolithography^31^, pre-polymer extrusion^32–38^ or molding^39^. Artificial vessels have also been formed by the direct removal of diverse hydrogel material such as silk fibroin and PEG hydrogels using laser photo-ablation^40–44^ including in the presence of cells^45–47^.

Despite the versatility of these vessel templating approaches, the minimal diameter of perfusable engineered vessels reported thus far has been limited to 150µm^33, 39^ and, in parallel with strategies based on extrinsic angiogenesis in organ-on-chip implementations^48^, the size of generated tissues has been limited to 400-500µm in at least one dimension^49–53^. Because of their small size, engineered tissues which have been implemented thus far do not preserve a physiologically relevant signaling context within the tissue, nor do they develop to a level of complexity comparable to in vivo organs.

### Photo-polymerizable non-swelling hydrogels enable 3D soft microfluidics

In order to vascularize tissue at large scale, we hypothesized that a microfluidic approach which could bridge the capillary to tissue scale would be necessary to enable the perfusion of thick three-dimensional tissue constructs. We therefore designed a dense, regularly spaced capillary network whose tubular walls were made of a hydrogel allowing diffusion (**Fig. 1a**). Tissues growing within this soft grid-like hydrated network would interface with an external perfusion pumping system, circulating cell culture medium throughout the volume of the tissue (**Fig. 1a** and **Supplementary Fig. 1**). We developed an approach whereby the perfusable grid would be printed directly on a hard plastic base (**Fig. 1a**), thereby forming a tight seal. This base, which contains perfusion holes, would then be incorporated into a perfusion chip linked to a peristaltic pump circulating cell culture medium (**Supplementary Fig.1**)

**Figure 1.**
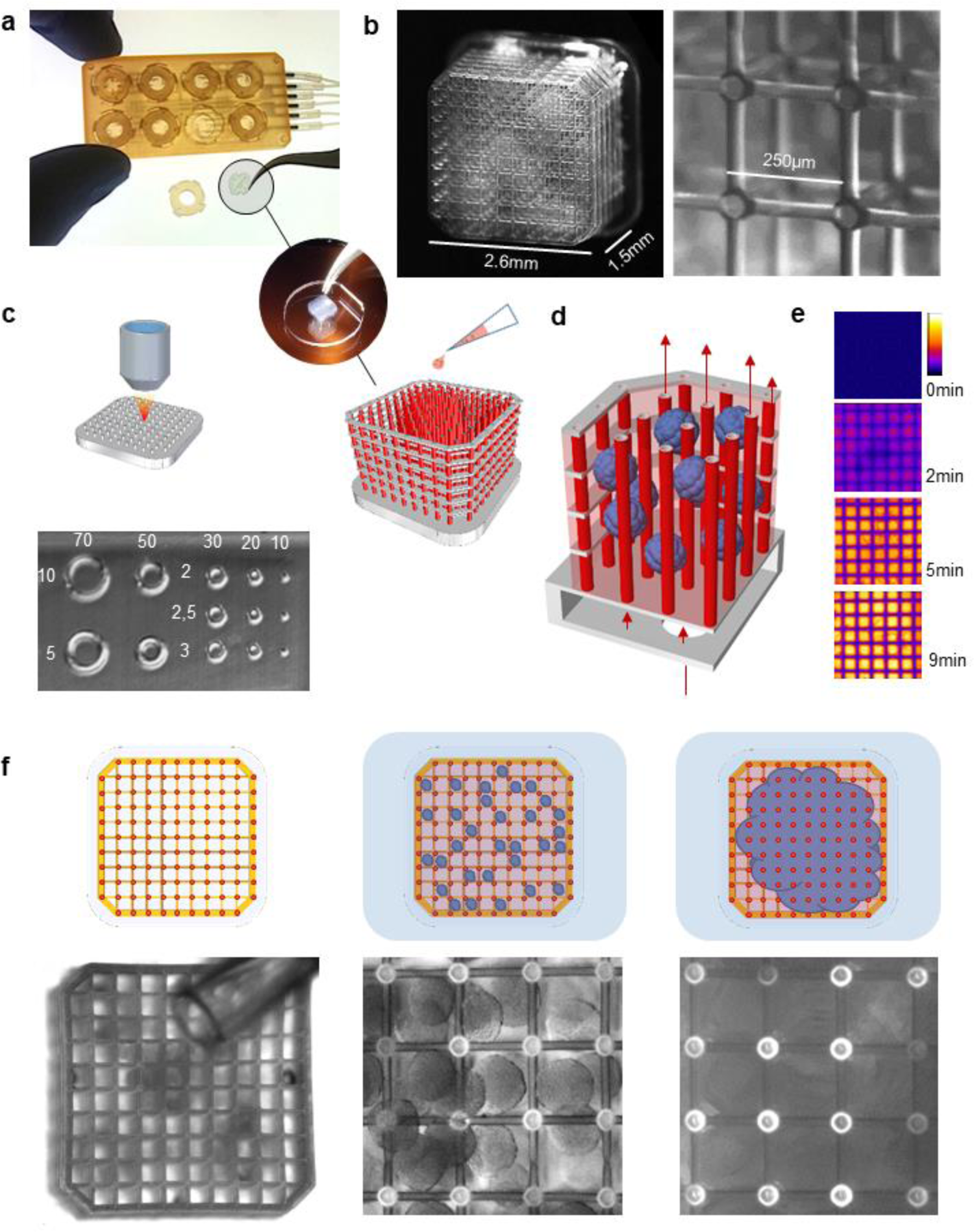
On-chip micro-vascularization enabled by soft microfluidics. (**a**) 3D printed tissue culture chip designed for eight multiplexed soft 3D soft microfluidic capillary grids. Inset: microfluidic capillary grid fabricated on a plastic baseplate and diagram of working principle. (**b**) of microfluidic capillary grid (left) and the close up image of the same grid showing individual hydrogel capillaries (right). (**c**) Microvessel 3D printing by 3D printing of microfluidic grid using high-resolution 2-photon stereo-lithography with non-swelling photo-polymerizable hydrogel precursors enables reproduction of features as small as 10µm. The top view photograph demonstrates an array of cylinders of various outer diameter in micrometers (top row) and wall thickness in micrometers (columns). (**d**) CAD image of microfluidic grid (left) with capillaries shown in red and the structural components shown in grey. The fence at the circumference of the structure makes up a “basket” that can be filled (right) with cell aggregates (spheroids) which merge and produce a solid tissue incorporating hydrogel capillaries. (**e**) Hydrogel capillaries are readily permeable. 25µM of Fluorescein, perfused through the microfluidic grid, embedded in Matrigel, passes across capillary walls and saturates the gel within several minutes. (**f**) Process of tissue generation. Photograph of an empty capillary grid with the tip of 200µl pipette tip visible above the grid (left), close up image of the greed seeded with hIPSC spheroids (middle) manually dispensed from the pipette shown on the left image, image of the spheroids fused into a solid tissue after 24-36 hours of culturing on chip. The top row schematically represents the same process, where spheroids and the resulting tissue are shown in blue.

An important requirement of this platform was the need to have capillary-like tubing at scales of a few µm in diameter and thickness, while perfusing across a large, multi-mm^3^ three-dimensional space. The geometrical complexity of the design, properties of the biopolymer and fabrication scale ranging from 10µm to 2000µm featured by our design made two-photon laser scanning photo-polymerization the ideal technology for this purpose. Initial 3D prints with the 2-photon Nanoscribe printer using commonly used photopolymerizable materials, including gelatin and PEG diacrylate, resulted in significant swelling of the material upon polymerization and hydration, which disrupted the seal between the soft microfluidic grid and the rigid plastic plate, and generated mismatched tubular segments (**Supplementary Fig. 2**).

To overcome this post-printing distortion, we developed a custom formulated hydrophilic photo-polymer based on polyethylene glycol diacrylate (PEGDA). While PEGDA exhibits cell-repelling surface properties, the polymer surface could be rendered cell-binding by the addition of the photocrosslinker pentaerythritol triacrylate (PETA)^54^. We reasoned that the addition of a significant amount of PETA as a crosslinker would increase the toughness of the polymer, and balanced with the addition of an inert “filler” component (Triton-X 100) would on the other hand retain sufficient porosity of the polymer to enable rapid diffusion.

The combination of 2-photon printing with this non-swelling hydrogel material allowed printing of a variety of microfluidic grids, ranging in size from 1.2 x 1.2 x 1.2mm up to 6.5 x 6.5 x 5mm (**Fig. 1b** and **Supplementary Fig. 3**) with vessel diameters from 10µm to >70µm and vessel wall thickness from 2µm to 10µm (**Fig. 1c**). Importantly, the printing with our novel formulation resin resulted in a 1:1 fidelity between the generated CAD geometry and the printed parts, thereby ensuring no distortion and a tight seal (**Supplementary Fig. 2**).The standard size used for most of the subsequent experiments was 2.6mm x 2.6mm x 1.5mm, with an inter-vessel distance of 250µm (**Fig. 1b**). These grids could be incorporated into a multi-plexed perfusion chip allowing up to 8 grids to be perfused simultaneously (**Fig. 1a**). Single cells or organoids smaller than the 250µm inter-capillary distance, previously mixed within a liquid hydrogel (eg. Matrigel) precursor solution, could be seeded into the platform, yielding a “gel-in-gel” 3-dimensional construct (**Fig. 1d**). Vessel permeability to water-soluble molecules within the chips overlayed with Matrigel was verified using fluorescein, with diffusion throughout the three-dimensional space seen in less than 10 minutes (**Fig. 1e**).

To perform a biological proof of concept experiment, we generated hundreds of organoids of less than 200 µm diameter by aggregating human pluripotent stem cells (hPSCs) in microwells in pluripotency medium over 24 hours. We collected these aggregates in cold liquid Matrgiel, which were then pipetted into the grids. Initial seeding demonstrated that these aggregates filled the grids and, over a period of 8 days of growth and neural differentiation, merged and filled the whole volume (**Fig. 1f** and **Supplementary Fig. 4**).

### scRNAseq reveals changes in differentiation, hypoxia, cell cycle regulation and differentiation upon perfusion

To assess how perfusion affected cellular processes and differentiation in large-scale in-vitro tissue, we dissociated cells from tissue constructs in a perfused and a non-perfused grid, as well as from organoids in conventional suspension culture after the 8-day culture period, and performed single cell RNA sequencing. Graph-based clustering and Uniform Manifold Approximation and Projection (UMAP) dimensionality reduction technique on the 8,625 total cells retained after QC revealed significant transcriptomic differences between the tissue constructs (perfused and non-perfused) and the control organoids as evidenced by largely separated clustering of these cell populations (**Fig. 2a**). Correlation analysis using the 100 most differentially expressed marker genes revealed the most difference between perfused tissue and conventional organoid culture, with non-perfused tissue sharing gene expression profiles with the other two conditions (**Supplementary Fig. 5**). Differential gene expression analysis was then used to annotate eight clusters, which were largely differentiated by fate, as well as by metabolic, hypoxia and cell cycle regulation (**Fig. 2b, Fig. 2d, Supplementary Fig. 6, Supplementary Fig. 7**). Cells from control organoids were found in clusters with low, medium and high glycolytic processes marked by a pluripotent identity, including highly expressed markers such as *NANOG* and *OCT4* (*POUF5F1*). Both perfused and non-perfused tissues expressed varying degrees of hypoxia, proliferation, mitochondrial gene expression and neuroepithelial markers. The non-perfused tissue made up the largest proportion of the hypoxic and stressed clusters (*HIF1α*, *FOS*) while the perfused tissue represented the majority of the neuroepithelial cluster (*PAX6*).

**Figure 2.**
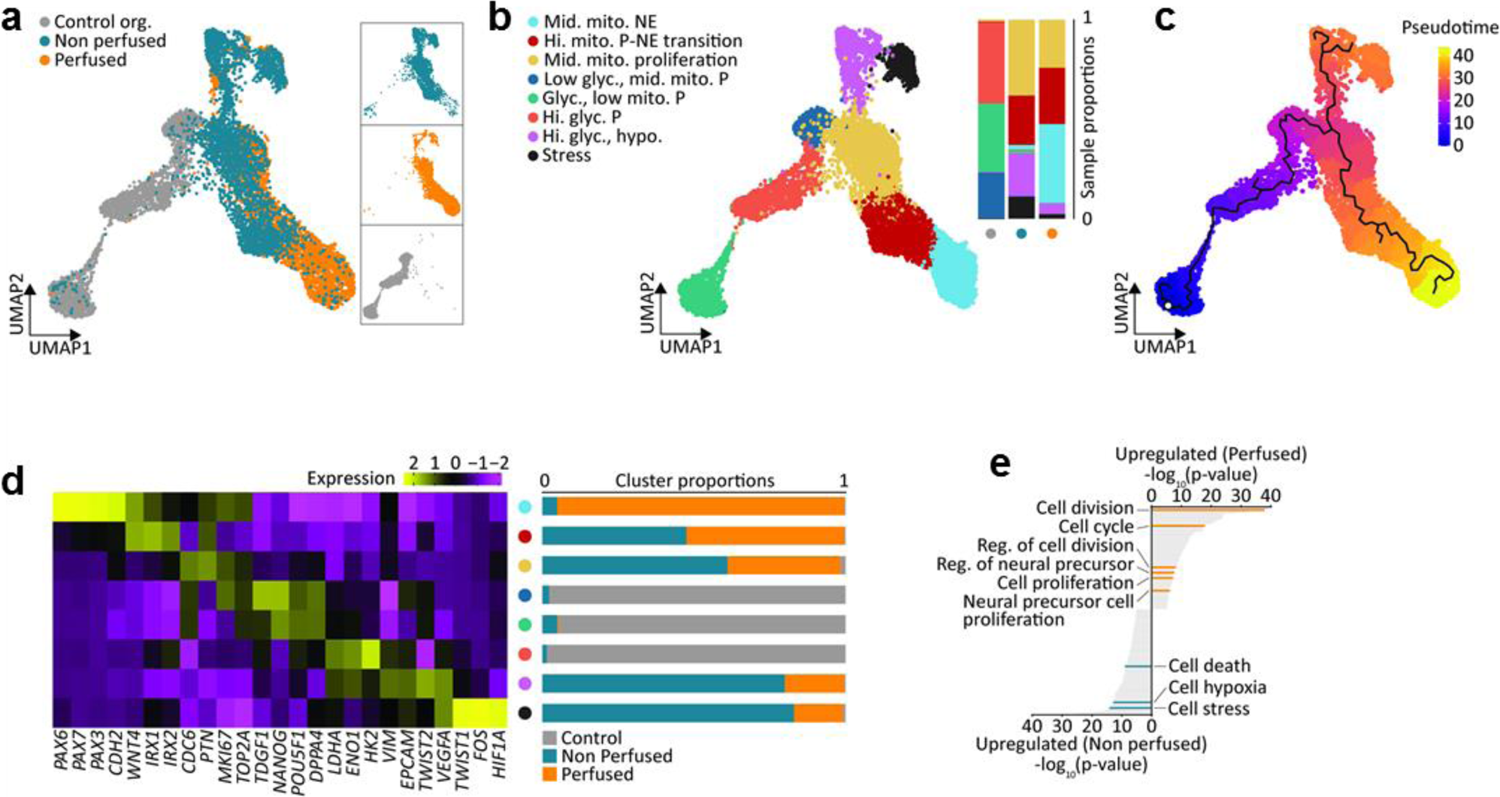
Transcriptomic changes upon perfusion. (**a**) Combined dataset UMAP (control, non-perfused and perfused samples). (**b**) Combined dataset UMAP with neuroepithelial cells (NE), pluripotent-neuroepithelial transitioning cells (P-NE), proliferating cells with medium mitochondrial (mito.) content, pluripotent cells (P) with low glycolysis (glyc.) and medium mitochondrial content, glycolytic pluripotent cells with low mitochondrial content, highly glycolytic pluripotent cells, a highly glycolytic hypoxic (hypo.) identity and a stressed cluster. (**c**) Pseudotime trajectory on combined dataset UMAP. (**d**) Cluster specific expressions of selected marker genes. Expression values are normalized and centered. Sample fractions for each identified cluster. (**e**) GO enrichment analysis for key processes upregulated in perfused and non-perfused samples.

In particular, control organoids had lower expression of mitochondrial genes (**Supplementary Fig. 7, Supplementary Fig. 8**) compared to the tissue constructs. This is in line with previous reports of lower mitochondrial activity in human embryonic stem cells that increase upon differentiation to fit the energy needs of resultant cell identities^55^. Indeed, differentiation towards the neuroepithelial fates from pluripotent cells was characterized by an increase in mitochondrial activity, and a simultaneous decrease in glycolysis (**Supplementary Fig. 8**). These results suggest a metabolic switch from anaerobic glycolysis to oxidative phosphorylation in the tissue constructs, as has been reported to occur during cell differentiation after exit from pluripotency^56, 57^. Such transitions, which were additionally evidenced in pseudotime analysis (**Fig. 2c**) distinguished the control organoids from the tissue constructs in the microfluidic grids, and further characterized differences between non-perfused and perfused tissues. The perfused tissue was also characterized by the expression of the neuroepithelial markers *PAX6*, *PAX7*, *PAX3*, and *CDH2*, while sharing *IRX1* and *IRX2* markers with non-perfused tissue (**Fig. 2d**). Moreover, while non-perfused and perfused tissues represented similar proportions of the pluripotent proliferating cluster, the perfused tissue expressed the highest proliferation markers such as *CDC6* (**Supplementary Fig. 7**). Additionally, the high expression of proliferation markers such *MKi67* and *TOP2A* in the neuroepithelial cluster suggested the retained proliferative capacity of perfused tissues after neuroepithelial differentiation, while maintaining overall minimal hypoxic/stress response markers *VEGF*, *FOS*, *HIF1a* (**Supplementary Fig. 8**). By contrast, non-perfused tissue was distinguished by significant presence of pluripotency (*NANOG*, *POU5F1*/*OCT-4*) (**Fig. 2d**) and hypoxia/stress markers (**Fig. 2d, Supplementary Fig. 8**). The control organoids, on the other hand, were mostly represented by pluripotent and hypoxic cells with complete absence of neuroepithelial identity (**Fig. 2d** and **Supplementary Fig. 8**), likely due to the short time in neural differentiation conditions.

To investigate transcriptional changes and associated cellular processes upon perfusion in a systematic manner, we performed differential gene expression analysis between perfused and non-perfused tissue constructs followed by Gene Ontology (GO) enrichment analysis (**Fig. 2e**). This analysis confirmed up-regulation of processes related to cell division and proliferation in the perfused sample, as well as regulation of neural precursors and neural precursor cell proliferation. By contrast, cell stress, hypoxia and cellular death processes were upregulated in non-perfused samples. Taken together our analysis of the transcriptomic data suggested that perfusion of large tissue constructs dramatically decreased apoptosis and hypoxia, and accelerated the process of neural differentiation.

### Perfusion rescues hypoxia and necrotic core in large tissue constructs

To further investigate how perfusion modulates the spatial distribution of hypoxia and apoptosis, we imaged the constructs in bright field microscopy and sectioned whole samples transversely, perpendicular to the direction of grid perfusion, followed by immunohistochemistry (**Fig. 3a** and associated quantification in **Fig. 3b**). Control organoids demonstrated a characteristic dark dense tissue core with occasional lighter void-like structures, surrounded by more translucent peripheral tissue. In non-perfused samples, tissue growth was restricted to the internal volume of the microfluidic grid, with generally dense central tissue interspersed with patchy lighter areas. Strikingly, perfused tissue covered the entire volume of the grid in a uniformly dense manner, with bulging epithelial outgrowths characteristic of cerebral organoids at the periphery. These observations suggested that cell proliferation was much higher in the perfused grids, compared to the non-perfused samples. To confirm this observation quantitatively, we dissociated the tissue constructs into single cells, stained with calcein-AM and ethidium homodimer to label live and dead cells respectively, followed by quantitative flow cytometry (**Fig. 3b** and **Fig. 3c**). Our analysis revealed a 5-fold difference in total cell number in the perfused tissue constructs over the non-perfused ones, while the proportion of live cells was similar (90.9±5.1% in perfused vs 89.2±4.4% in non-perfused tissue) (**Fig. 3a** and **Fig. 3b**).

**Figure 3.**
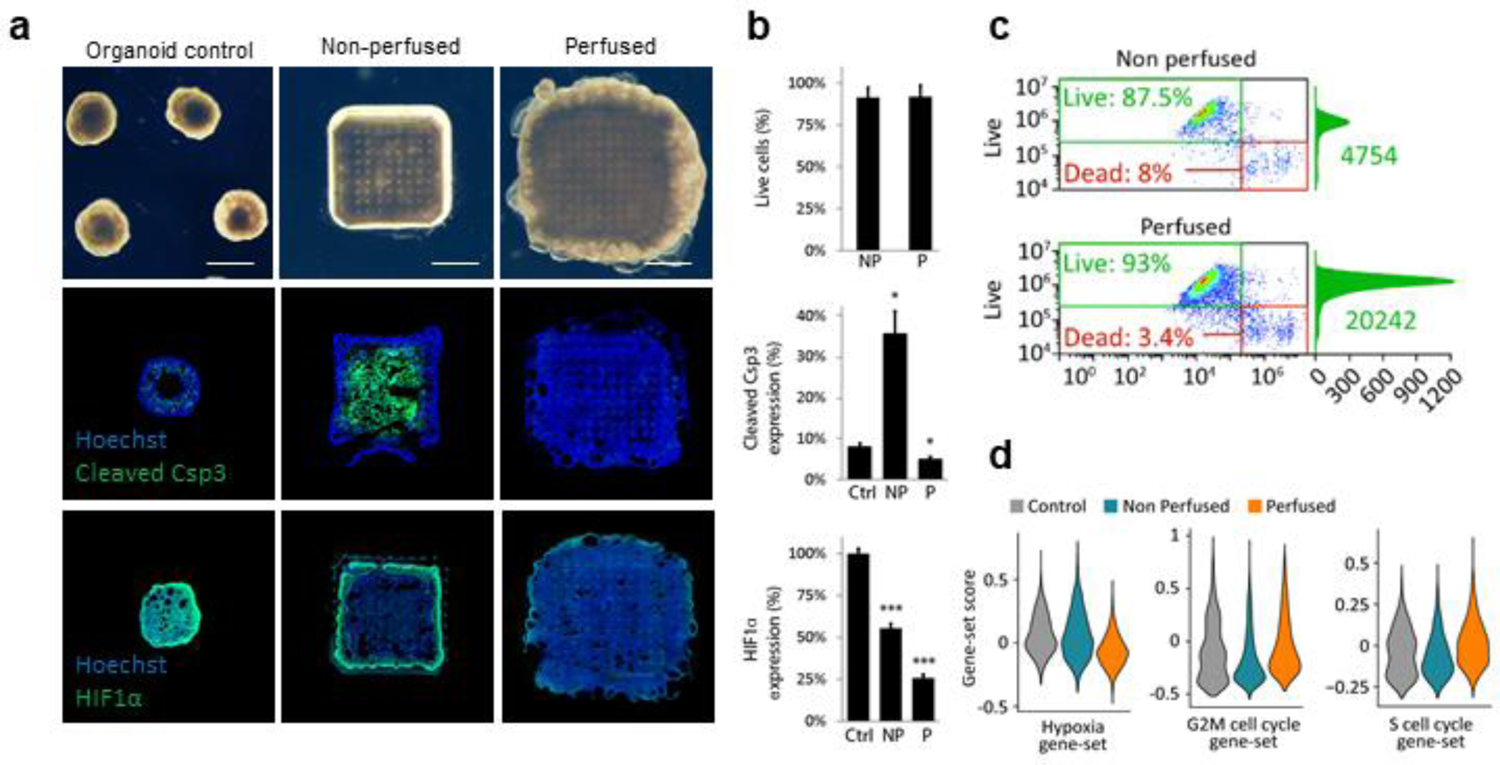
Changes in proliferation mediated by hypoxia. **a**) Representative experimental results demonstrating differences in proliferation and viability between standard organoid culture(left column) and the tissue constructs without (middle column) and with perfusion (right column). Top row: bright field images of organoids, non-perfused and perfused constructs correspondingly. Middle row: immunofluorescent images of apoptotic marker cleaved Caspase 3 (green) expressed in the three conditions. Bottom row: immunofluorescent images of hypoxia marker HIF1a (green). Hoechst staining of nuclei shown in blue. The images represent transverse cross-sections of the tissue constructs. (**b**) Top: average proportion of the number of live cells in perfused and non-perfused constructs quantified by flow cytometry (90.9±5.1% in perfused vs 89.2±4.4% in non-perfused tissue, n.s., n=4). Middle: average expression of cleaved Caspase 3 in control organoids (8±1% area, n=7), non-perfused(35.7±5%, p_Control_=0.02, n=4) and perfused tissue constructs (5.1±1% of total area, p_Control_=0.02, n=9). Bottom: average expression of hypoxia marker HIF-1a in control organoids (100±3%, n=4), non-perfused (55±3%, p_control_=0.0003, n=3) and perfused constructs (26±2%, p_control_=0.0004, n=2). Control organoids, non-perfused and perfused constructs are denoted as Ctrl, NP and P correspondingly, data are represented as mean ± SEM, asterisks (*) denote statistical significance of the difference between control organoids and the tissue constructs (unpaired two-tailed Student’s *t*-test, 95% confidence interval). (**c**) Representative fluo-cytometry data, demonstrating the difference in total number of cells between perfused and non-perfused constructs. (**d**) Distribution of gene expression levels across the single cell population for hypoxia (left), G2M cell cycle (middle) and S cell cycle (right) associated gene sets. Scale bar: 1mm

To determine whether these changes in viability and proliferation were due to apoptosis, we stained sections of the grids for cleaved Caspase 3, an active form of the Caspase-3 enzyme responsible for the degradation of multiple cellular proteins and ultimately for cell fragmentation into apoptotic bodies. Upon sectioning, the control organoids were empty in the center, suggesting, as has previously been reported^58^, the loss of apoptotic cells during the sectioning process. Closer to this inner core, signs of apoptosis were evident (8±1% of total area), (**Fig. 3a,b**), consistent with the large size of these organoids. Tissue in the non-perfused grids exhibited a clear inner core of apoptotic cellular fragments (35.7±5% of total section area) (**Fig. 3a**) along with empty regions completely lacking cells, with some analogous features to control organoids. Strikingly, nearly the entire perfused tissue did not show signs of cleaved Caspase 3 (5.1±1% of total area), indicating that perfusion successfully prevented cell apoptosis throughout the course of differentiation.

To verify whether hypoxia could be involved in initiating the observed apoptosis in the non-perfused samples, we next stained for HIF1α, a heterodimer protein complex playing a key role in oxygen homeostasis^59, 60^. The rapid buildup of HIF-1α in low-oxygen conditions is known to trigger a hypoxic response ultimately leading to apoptosis^61, 62^. In line with scRNA analysis data (**Fig. 3d**), high HIF1α expression levels were detected in the control organoids (100±3% mean fluorescent intensity), as well as in the non-perfused samples (55±3% of organoid control). Conversely, low levels of HIF1α expression were observed in the perfused samples (26±2% of organoid control) (**Fig. 3a**), associated with a higher proportion of cycling G2, M and S phase cells (**Fig. 3d**).

The patterns of HIF-1α and cleaved Caspase3 expression in non-perfused tissue sections therefore suggest that in these samples, cells in the center of the construct were in a transient hypoxic state ultimately leading to apoptotic cell death. This transition from hypoxic to apoptotic cell state continued until the volume of the tissue was small enough to allow sufficient oxygen supply by a passive diffusion, with the accumulation of apoptotic bodies and cellular debris preserving the cleaved Caspase 3 expression in the bulk of the non-perfused tissue (**Supplementary Fig. 9**). Conversely, this phenomenon was completely absent in the perfused samples, clearly confirming that thick tissues at multi-mm^3^ scale could be grown with high viability, and with little to no apoptosis or hypoxia within this platform.

### Accelerated neural differentiation in perfused tissue constructs

Our scRNAseq data suggested that perfusion not only improved proliferation, prevented apoptosis and hypoxia, but could also direct fate specification. To confirm these findings, we analyzed specific markers of pluripotency and neural differentiation by immunohistochemistry (**Fig. 3a** and associated image quantification in **Fig. 3b**). NANOG, a canonical marker of pluripotency was abundantly expressed in organoids (73.6±8% of cells) and significantly expressed in non-perfused constructs (16.1±2%) at the protein level, but was completely missing in perfused samples. Conversely, PAX6, the earliest marker of neural differentiation was clearly evidenced in the perfused samples (21.5±2%), but not in organoid controls and non-perfused samples. These results were in line with the scRNAseq data, which indicated that *NANOG*-expressing cells were present in the control and non-perfused samples, while *PAX6* was largely expressed in the perfused sample (**Fig. 4c**). The transition of stem cells from pluripotency to neural identity is regulated by a loss of cellular expression of E-CADHERIN (CDH1) and a gain of N-CADHERIN (CDH2) expression which drives neural differentiation by inhibiting FGF-mediated pathways^63^. In order to assess whether perfusion enhanced this transition, we stained for E-CADHERIN to evidence the epithelial state associated with pluripotency, and for N-CADHERIN to confirm the transition to early neuro-epithelial identity. Perfused samples were indeed largely N-CADHERIN positive (55.7±9% vs 7.6±1% in non-perfused samples), while more cells in the non-perfused samples maintained E-CADHERIN+ identity (26.6±7% vs 4.5±2% in perfused samples). These results were confirmed by the scRNA data (**Fig. 4c**), indicating the predominant expression of N-CADHERIN in cells from perfused tissue, and E-CADHERIN in non-perfused and control tissue. Taken together, these results demonstrate that perfusion rapidly accelerates the transition from pluripotency to early neuroepithelial identity, while cells which lack perfusion remain in a state of pluripotency.

**Figure 4.**
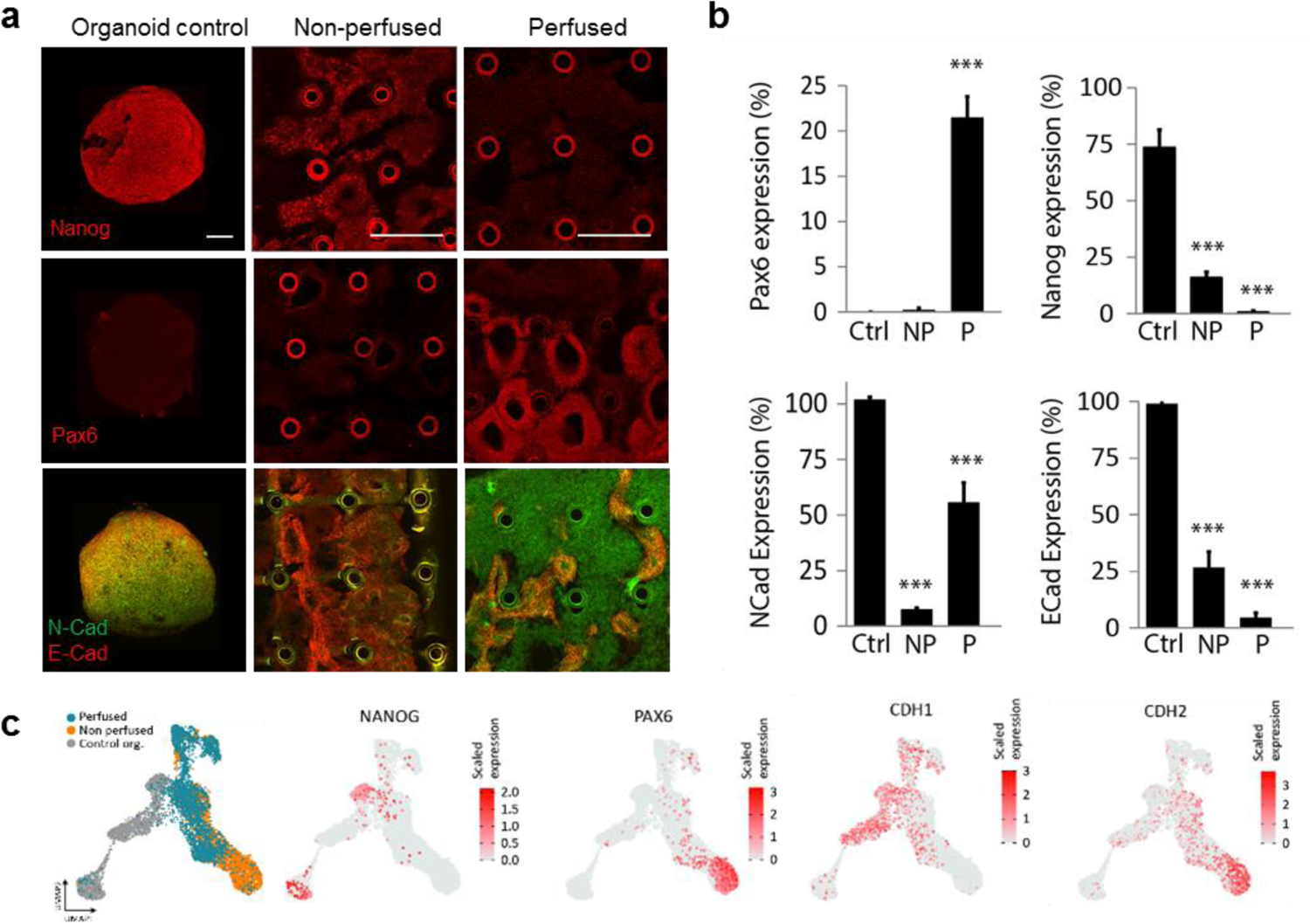
Rapid neural differentiation in perfused systems. (**a**) Representative experimental results demonstrating the result of 2 days of neuronal induction in control organoids(left column), non-perfused (middle column) and perfused (right column) tissue constructs. Top, middle and bottom row: immunofluorescent images of stem cell marker NANOG, early neural marker PAX6 and cell adhesion proteins N-Cadherin (green) and E-Cadherin (red) correspondingly. The images represent transverse cross-sections of the tissue constructs. In most cases, the cross-sections of capillaries are visible as circular structures (marked by triangles) (**b**) Average expression of PAX6(Ctrl: Not detected; NP: 0.2%±0.2%, p_Control_= 0.35(n.s.), n=9; P: 21.5±2%, p_Control_=0.00002, n=12), NANOG(Ctrl: 73.6±8%, n=9; NP: 16.1±2%, p_Control_=0.00007, n=11; P: not detected), NCad(CDH2) (Ctrl: 100±1%, n=8; NP: 7.6±1%, p_Control_=4e-13, n=3; P: 55.7±9%(p_Control_=0.009, n=5) and ECad(CDH1) (Ctrl: 99±2%, n=12; NP: 26.6±7%, p_Control_=0.0001, n=6; P: 4.5±2%, p_Control_=2e-10, n=5) markers in control organoids (Ctrl), non-perfused(NP) and perfused(P) constructs. Data are represented as mean ± SEM, asterisks (*) denote statistical significance of the difference between control organoids and the tissue constructs (unpaired two-tailed Student’s *t*-test, 95% confidence interval) (**c**) UMAP plot of the combined dataset showing the localization of cells from control organoids, non-perfused and perfused constructs in the UMAP space. (**d**) UMAP plot of the combined dataset highlighting locations of PAX6, NANOG, NCad (CDH2) and ECad(CDH1) expressing cells in the UMAP space. Scale bar: 250µm

### Long-term perfusion of liver tissue constructs

To demonstrate the versatility of this platform, we went on to assess the differentiation of PSC-derived liver progenitors in perfused grids. hPSC were differentiated for 8 days in conventional 2D culture, followed by spheroid generation, seeding into the chip and perfusion for an additional 32 days. As was the case with neural differentiation, cells merged over time into a continuous tissue (**Supplementary Fig. 10**), with histological analysis identifying tightly packed cells with an eosinophilic and clear vacuolated cytoplasm reminiscent of hepatocytes, with no evidence of apoptotic bodies or necrosis throughout the tissue (**Fig. 5a**). The expression of many hepatocyte-specific genes was upregulated in perfused tissue constructs compared to both control hepatic organoids and 2D hepatocyte differentiation, including hepatocyte nuclear factor 6 (HNF6), Na^+^/taurocholate co-transporting polypeptide (*NTCP*), albumin (ALB), alpha1-antitrypsin (AAT) and two major cytochrome *P450* enzymes CYP2C9 and CYP3A4 (**Fig. 5f**). The presence of CYP3A4 as well as of the hepatocyte progenitor marker AFP were confirmed at the protein level in the perfused sample by immunohistochemistry (**Fig. 5b**). Interestingly, the two major gluconeogenesis enzymes phosphoenolpyruvate carboxykinase (PEPCK) and glucose 6-phosphatase alpha (G6PC) were also expressed at higher levels in the perfused constructs, compared to control organoids (1.6- and 1.3-fold higher expression, respectively) (**Fig. 5f**).

**Figure 5.**
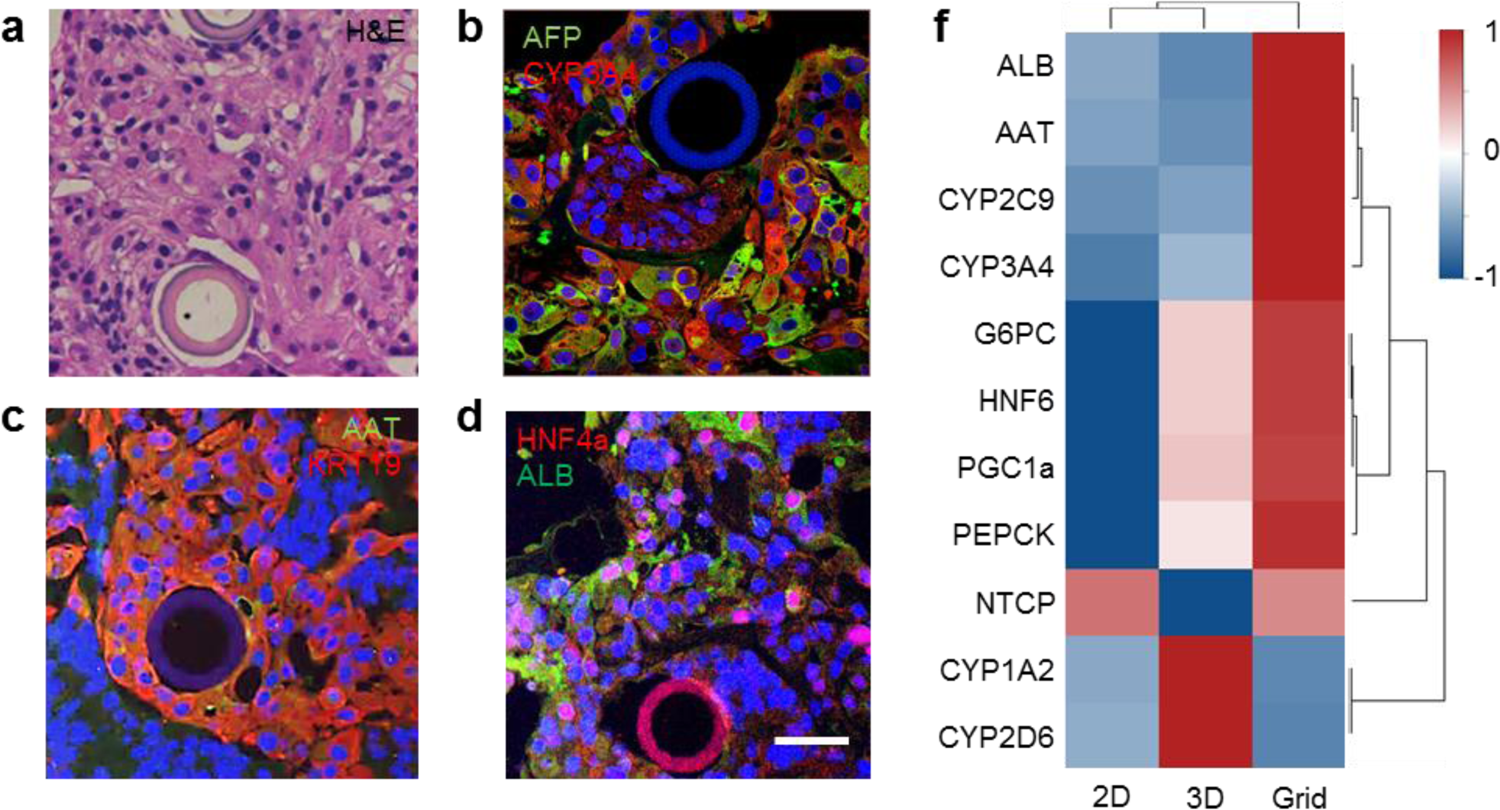
Functional improvement in perfused liver microtissues. (**a**) Hematoxylin and eosin of a transverse section of liver tissue construct with hepaptocyte-like morphology and no visible indication of apoptosis. (**b**) Immunofluorescence images showing presence of AFP and CYP3A4, (**c**) Alpha-1 antitrypsin (AAT) and Cytokeratin 19 (KRT19), and (**d**) basic hepatic markers albumin (ALB) and hepatic nuclear factor HNF4α expression (**f**) Heatmap representation of fold change gene expression levels normalized to control 2D cell culture and compared to standard hepatic organoids and perfused liver constructs; rows are centered and scaled. Gene expression data for hepatic 2D culture and hepatic organoids are from Kumar et al^74^. Scale bar: 50µm

We next assessed whether non-parenchymal cells in the developing liver, such as cytokeratin 19 (*KRT19*)-expressing cholangiocytes, which contribute to bile secretion and hepatocyte survival^64^, were also present in our perfused culture system. These cells are generated *in-vivo* from hepatoblasts surrounding the portal veins, while hepatoblasts located away from portal vein areas differentiate into hepatocytes^65^. We observed a similar localization pattern of KRT19+ cells, with such cells tightly surrounding every vessel in the microfluidic grid (**Fig. 5c**), while the hepatocytic markers HNF4α^66^ was expressed in cells scattered in the inter-vessel space (**Fig. 5d**). Additionally, the functional production of albumin (ALB) was confirmed by staining (**Fig. 5d**).

Taken together, our results confirmed the feasibility of using this synthetic micro-vascularization approach as a generic strategy to building large, viable, perfused in-vitro tissues.

## Discussion

Here, we demonstrated for the first time an integrated3D culture platform which provides a physiologically relevant micro-perfusion for engineered tissues, resulting in enhanced tissue growth and differentiation compared to previously reported in-vitro tissue vascularization strategies.

We showed that microvascular networks could be created using 2-photon hydrogel polymerization, demonstrating that this technology can be used to achieve micro-vasculature with previously unreported accuracy, resolution and scale. A significant limitation of current photo-polymerizable hydrogel materials is the significant swelling of the material, which prevents the robust, leak-free interface between printed structures and microfluidic perfusion systems. Our development of a non-swelling photo-polymerizable material formulation was a critical component to overcome this limitation, and enabled the printing of 3D soft microfluidic systems which could be reliably perfused over multiple weeks. The exchange of nutrients and oxygen as well as the removal of waste products was achieved via simple diffusion as the printed vascular network is permeable to water-soluble molecules and gases. The direct fabrication of capillaries of a defined topology delivers an unprecedented control over tissue perfusion parameters, and the design of the vascular network is highly flexible and can be adapted to more complex geometrical and structural requirements. To provide a complete extracellular milieu with structural 3D support to the growing engineered tissue, the space around the perfused vessels was filled with hydrogel. In the experiments presented here, the hydrogel component consisted of the commonly used proteinaceous matrix Matrigel, however the platform can also readily accommodate without any additional modifications the use of other naturally derived matrices such as collagen, as well as synthetic artificial extracellular matrices such as poly(ethylene) glycol PEG^67^ or alginate^68^. This technology provides a reliable fluidic coupling between the microfluidic grid and the host perfusion device, such that a continuous peristaltic pump driven perfusion is possible. By integrating these printed microfluidic grids into a perfusion system, we were able to demonstrate that that large-scale (>15mm^3^) fully perfused neural and liver tissues could be generated with this platform.

Our experiments with neural tissue demonstrate that the differentiation trajectory of cells in this perfused system is significantly enhanced. While control organoids remained largely in a pluripotent state, scRNAseq analysis revealed that perfused tissue rapidly differentiated towards the neural fate, together with a switch from glycolysis to oxidative phosphorylation. Imaging and flow cytometry confirmed that this tissue was highly viable, and immunohistochemistry showed markers of neural differentiation, which were absent in the non-perfused sample, as well as hallmarks of epithelial to mesenchymal transition. This was underscored by our observations that the lack of active micro-perfusion of the engineered tissue construct triggered a stress response in the cells within the inner core of the tissue. The accelerated differentiation of tissues upon perfusion is thought to be due not only to increased availability of nutrients and oxygen but also to the rapid diffusion of differentiation factors within the tissue via the tightly spaced capillary network.

The possibility of long-term perfusion within this platform was demonstrated by the maintenance of viable engineered liver tissue demonstrating enhanced phenotypic and functional features compared to standard 2D and 3D organoid culture. As cells in our current capillary design were not sufficiently distant from the source of oxygen to generate an oxygen gradient, we did not observe a clear spatial segregation of hepatocytic markers, known as zonation, in the perfused liver tissue, however this feature could be engineered by wider inter-vessel distance.

Overall, the 3D soft microfluidic technology presented here overcomes one of the major challenge in engineering tissues and organoids: the lack of tissue perfusion from the initiation of tissue growth, and enables the generation of large engineered tissues which are vascularized from within and their maintenance over long periods of time. In applications such as disease modeling and drug development, such a highly defined synthetic perfusion system would be beneficial in avoiding the complexity and variability introduced by exogenous angiogenesis-driven vascularization. While the current implementation of the platform does not recapitulate features of in vivo vascular networks such as adaptive vascular remodeling or selective blood brain barrier interactions, it addresses the major problem of oxygen, nutrient and growth factor and small molecule supply as well as of waste removal, allowing to generate viable tissues beyond currently available dimensions. The incorporating of endothelial vasculature with this synthetic capillaries could be implemented, where the hydrogel micro-vascularization could provide a temporary tissue support during the time required for angiogenesis-driven capillarization to establish a perfusable network. We expect that this approach is widely applicable in overcoming the current size limitations of bio-printed tissues and provides a technological foundation for the development of perfusable in vitro models of increased complexity and scale.

## Materials and Methods

### Human PSC culture

Human PSCs were cultured in Matrigel (356277, Becton Dickinson) coated 6 well plates up to 60-70% confluency. Passages were performed by a 3 min treatment of Dispase II (D4693, Sigma) at 37°C, followed by 2-3 PBS washings at RT. 1 mL of E8-Flex medium (A2858501, Thermo Scientific) was added and the colonies were scraped and gently pipetted 4-5 times through 1ml plastic tip to break the colonies. The colony suspension was then diluted at 1:5 ratio and plated to a Matrigel coated wells in 2 mL of E8Flex medium supplemented with 10 μM Rock inhibitor(Y-27632, Hellobio) for 24 h. The medium was then replaced by 2 mL of fresh E8-Flex medium and incubation was continued for 48 h, at which point the colonies usually reached 60-70% confluency and were ready for next passage.

### Human PSC-derived cerebral organoids and perfused cerebral tissue

We adapted the protocol by Lancaster et al.^69^ to our experimental conditions. Upon reaching a confluency of 60-70% hPSCs were dissociated by treatment the colonies with 250µl Accutase (A1110501, Gibco) for 7 min at 37°C and re-suspended in E8Flex medium containing 10 μM Rock inhibitor.

Organoids were generated in U-bottom 96-well plates (#351177, Falcon). The plates were rinsed with Anti-Adherence Solution (#07010, Stemcell Technologies) and cells were plated at 9000cells/well density. Plates were spun at 300rcf at RT and left in CO2 incubator for 24 hours for hPSC spheroid aggregation, after which culture medium was replaced by fresh E8-Flex medium, without Rock inhibitor and changed afterwards every 2 days. At day 2, the spheroids were embedded in growth factor reduced (GFR) Matrigel (354230, Becton Dickinson) and kept in 6-well plates, pre-rinsed with Anti Adherence Solution in CO2 incubator. At day 6, neuronal induction was started by replacing E8-Flex medium with DMEM/F12 medium (31330038, Gibco) containing 1% MEM-NEAA (11140035, Gibco), 1% Glutamax (35050038, Gibco), 1% Pen-Strep (15140122, Gibco), 0.5% N2 supplement (17502048, Gibco) and 1ug/ml Heparin (H3149, Sigma) (neural induction medium). Day 8 spheroids were used for characterization.

For the formation of perfused tissue, at day 0, we first generated micro-hPSC spheroids using 24-well Aggrewell plates (34411, Stemcell Technologies) following a protocol supplied by the manufacturer. Specifically, we seeded Aggrewells with hPSCs to obtain 350-400 cells per micro-well. After 24 hours, spheroids formed and the medium was replaced by fresh E8-Flex without Rock inhibitor. At day 2, the spheroids typically reached a diameter of 180-200um, and were harvested and resuspended in ice cold GFR Matrigel at a density of 3500 spheroids per 200-250ul GFR Matrigel. The grids were placed under stereomicroscope in a 35mm petri dish on ice and seeded with the spheroid suspension at final density of ∼1800 spheroids per grid. 200µl plastic tips pre-chilled on ice were used to dispense the suspension under stereomicroscope. Seeding was performed in several stages in order to allow dispensed spheroids to settle into the grids. After seeding, the grids were kept in a CO2 incubator for 40-50 min to allow Matrigel polymerization, after which the grids where placed in a perfusion chip and perfusion was started. E8-Flex/PenStrep medium was changed every 2 days (3-4ml per grid). At day 6, E8-Flex medium was replaced by neural induction medium for 2 days. At day 8, the grids were extracted and the tissue used for characterizations.

### Immunofluorescence analysis of cerebral tissue samples

Samples were fixed with 4% paraformaldehyde (158127, Sigma-Aldrich) for 24-36 hours at 4°C and washed by 3 incubations in PBS for 15-20 min at room temperature. Fixed tissue was sectioned either by embedding in low melting point agarose and sectioning on vibratome (Leica VT1000S) into 100-150µm sections or cryopreserved in OCT(6502, Thermo APD Consumables) overnight at 4°C, re-embedded into fresh OCT, frozen in isopropanol/dry ice slurry, cut into 50µm sections on cryotome (Leica CM1850) and affixed on SuperFrost Plus (10019419, Thermo Scientific) microscope glass slides.

Sections were incubated in a permeabilization and blocking solution of 0.3% Triton X (A4975, PanREAC AppliChem) and 3% BSA (A7906, Sigma) in PBS for 24 hours at 4°C. Primary antibodies were diluted in the permeabilization and blocking solution and applied to sections for 24h at 4°C, after which three PBS washes were performed over another 24h period. Secondary antibodies and Hoechst were also diluted in the permeabilization and blocking solution and applied to sections overnight at 4°C, followed by washing in PBS 3-4 times over another 24 hours. Antibodies used in this study are listed in Table 1. Stained agarose-embedded sections were stored in 2mM sodium azide solution in PBS. Cryosections were mounted in Fluoromount-G medium and stored at 4°C.

**Table 1.** List of antibodies used in this study

| Primary | Host | Dilution | Manufacturer |
| --- | --- | --- | --- |
| Pax6, monoclonal | Mouse | 1:200 | DSHB |
| Nanog | Goat | 1:200 | R&D Systems, AF1997 |
| ECad | Mouse | 1:500 | Abcam, ab76055 |
| NCad | Rat | 1:200 | DSHB |
| Cleaved Caspase3 | Rabbit | 1:400 | Cell Signaling Technology, 9661 |
| HIF1a | Rabbit | 1:500 | Abcam, ab51608 |
| HNF4α | Mouse | 1:200 | Abcam, ab41898 |
| Alpha-1-Antitrypsin | Rabbit | 1:200 | DAKO, A0012 (00092029) |
| PEPCK | Mouse | 1/1000 | Santa Cruz, sc-271204 |
| MRP2 | Mouse | 1/500 | Abcam, ab3373 |
| KRT19 | goat | 1/500 | Santa Cruz, sc-33120 |
| ALB | Rabbit | 1/500 | Abcam, ab207327 |

**Table 1. List of antibodies used in this study**
| Secondary |  |  |  |
| --- | --- | --- | --- |
| Hoechst |  |  | Sigma-Aldrich, 14533 |
| Anti-mouse Alexa 647 | Donkey | 1:500 | Invitrogen, A31571 |
| Anti-goat Alexa 647 | Donkey | 1:500 | Invitrogen, A21447 |
| Anti-rat Alexa 555 | Goat | 1:500 | Invitrogen, A21434 |
| Anti-goat Alexa 555 | Donkey | 1:500 | Invitrogen, A21432 |
| Anti-rabbit Alexa 555 | Donkey | 1:500 | Invitrogen, A31572 |
| Anti-mouse Alexa 488 | Donkey | 1:500 | Invitrogen, A11029 |

### Human PSC-derived perfused hepatic tissue

All liver differentiation experiments were performed with the H3CX hiPSC line previously generated ^70^. H3CX is a hIPSC line (Sigma 0028, Sigma-Aldrich) genetically engineered to overexpress 3 transcription factors HNF1A, FOXA3 and PROX1 upon Doxycycline induction, which allows for rapid generation of hepatocyte-like progeny. H3CX cells were expanded feeder-free on Matrigel (BD Biosciences)-coated plates in E8 or E8 Flex (Thermo Fisher Scientific). HC3X cells were differentiated towards HLCs as previously described^71^. Briefly, HC3X cells were dissociated to single cells using StemPro™Accutase™ Cell dissociation Reagent (Thermo Fisher Scientific) and plated on Matrigel-coated plates at ±8.75 × 10^4^ cells/cm^2^ in mTeSR medium (Stem Cell Technologies) supplemented with RevitaCell (Thermo Fisher Scientific). When cells reached 70-80% confluence, differentiation was performed during 40 days in liver differentiation medium (LDM) containing to comprise 500ml total volume: 285 ml of DMEM low glucose (Invitrogen 31885023), 200 ml of MCDB-201 solution in water (Sigma M-6770) adjusted to pH 7.2, 0.25× of Linoleic acid—Bovine serum albumin (LA-BSA, Sigma L-9530), 0.25× of Insulin–transferrin–selenium (ITS, Sigma I-3146), 50 U of Penicillin/Streptomycin (Invitrogen 15140122), 100 nM of l-ascorbic acid (Sigma A8960), 1 μM dexamethasone (Sigma D2915) and 50 μM of β-mercaptoethanol (Invitrogen 31350010). Differentiation medium was supplemented with 0.6% dimethylsulfoxide (DMSO) during the first 12 days of the culture. 2.0% DMSO and 3x concentrate of non-essential amino-acids (NEAA) was added to LDM medium between days 12-13, and from day 14 until the end of differentiation 20g/L glycine was added to LDM medium supplemented with NEAA. Differentiation was performed in presence of the following factors: day 0-1: 100ng/ml Activin-A and 50ng/ml Wnt3a, day 2-3: 100ng/ml Activin-A, day 4-7: 50ng/ml BMP4, day 8-11: 20ng/ml FGF1, and 20ng/ml HGF during the rest of differentiation. Doxycycline (5ug/ml) was applied from day 4 until the end of differentiation. All cytokines were purchased from Peprotech.

### RNA extraction and quantitative reverse-transcription PCR

RNA extraction was performed using TRIzol reagent (Invitrogen) following manufacturer’s instructions. At least 1µg of RNA was transcribed to cDNA using the Superscript III First-Strand synthesis (Invitrogen). Gene expression analysis was performed using the Platinum SYBR green qPCR supermix-UDG kit (Invitrogen) in a ViiA 7 Real-Time PCR instrument (Thermo Fisher Scientific). The sequences of all used RT-qPCR primers are listed in Table 2. The ribosomal protein L19 transcript (RPL19) was used as a housekeeping gene for normalization.

**Table 2.**
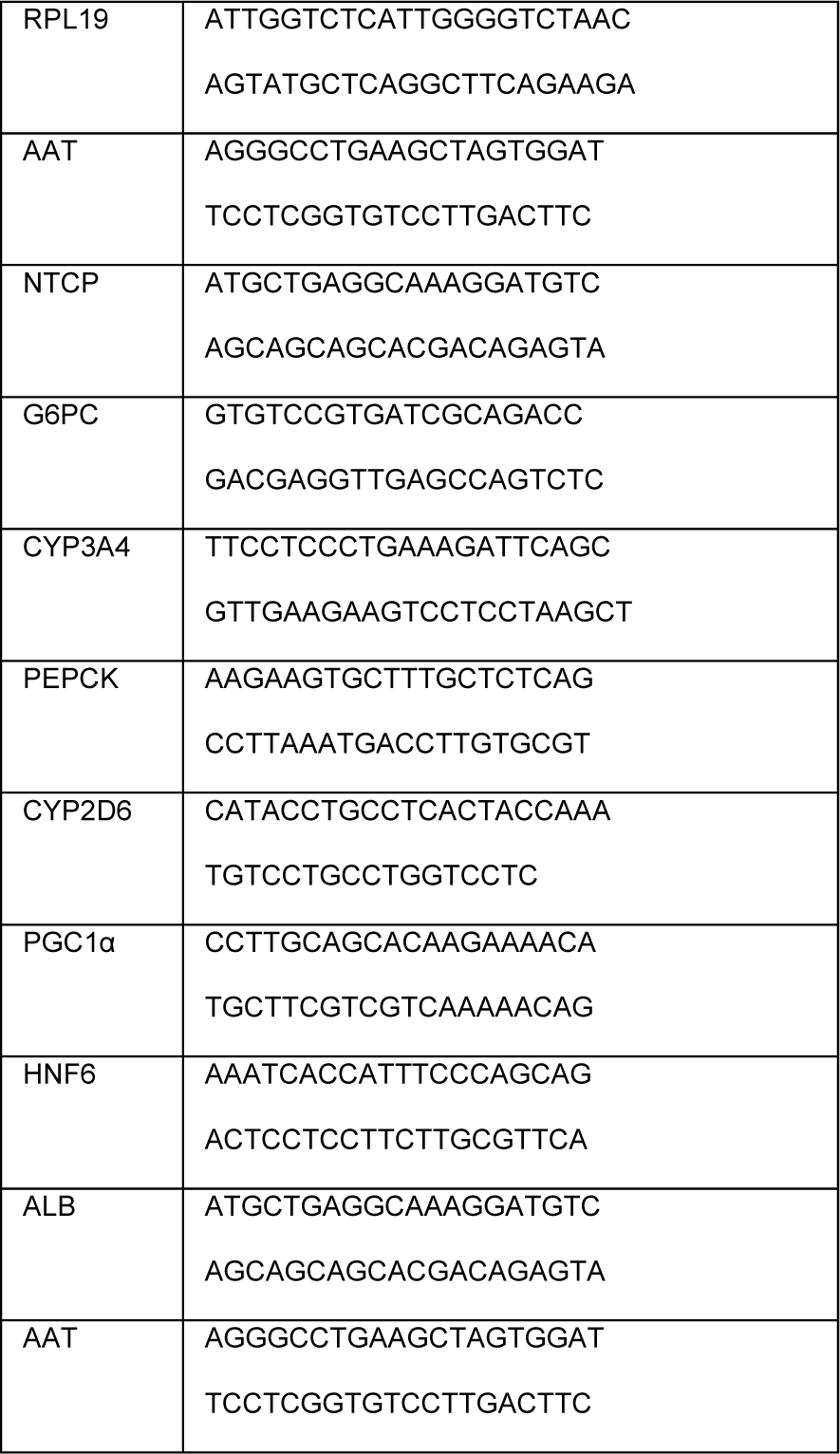
List of primers used in this study

### Histology

Samples were fixed with 4% (w/v) paraformaldehyde (PFA, Sigma-Aldrich) overnight at 4°C, washed 3 times with PBS and submerged in PBS-sodium azide (0.01% v/v) solution at 4°C until embedded in paraffin. Hydrogel sections (5 µm) were prepared using a microtome (Microm HM 360, Marshall Scientific.) For Hematoxylin and Eosin (H&E) staining, sections were treated with xylene solution to remove the paraffin, and gradually rehydrated in ethanol (100 to 70%, v/v). H&E staining was performed by submerging rehydrated hydrogel sections in Harris Hematoxylin solution, acid alcohol, bluing reagent and Eosin-Y solution by order. Stained samples were dehydrated with ascending alcohol series, washed in xylene solution, and mounted with DPX mountant (Sigma-Aldrich).

### Immunofluorescence analysis of liver tissue samples

Following deparaffinization in xylene and rehydration in descending alcohol series, heat-mediated antigen retrieval was performed by incubating hydrogel sections in Dako antigen retrieval solution (Dako) for 20 min at 98 °C. This step was followed by cell permeabilization with 0.01% (v/v) Triton-X (Sigma-Aldrich) solution in PBS, for 20 minutes. Samples were then incubated with 5% (v/v) Goat or Donkey Serum (Dako) for 30 min. Primary antibodies diluted in Dako antibody diluent solution, were incubated overnight at 4°C, followed by washing steps and incubation with Alexa-coupled secondary antibody (1:500) and Hoechst 33412 (1:500) solution for 1 hour at room temperature. Finally, samples were washed in PBS, and mounted with Vectashield antifade mounting medium (Vector Laboratories). Stained sections were imaged using laser scanning confocal microscope (LSM 880, Zeiss, Germany), and image processing were performed on ZEN Blue software (Zeiss, Germany).

### Image acquisition and analysis of cerebral tissue samples

Fluorescence images were obtained using a confocal microscope (Leica SP8 DIVE, Leica Microsystems) equipped with 10x NA0.4 dry objective. Acquisition parameters were kept constant for all samples obtained in the same experiment. Image processing was performed using Fiji/ImageJ (NIH) and custom-written ImageJ Java plugin. At the data preprocessing stage, image stacks were collapsed by maximum intensity projection. Resulting images were translated to 8-bit lookup table representation, manually cleaned from grid structure elements and thresholded by replacing all sub-threshold pixels with black (zero value) pixels. Lookup tables of the images were then remapped from [Threshold..255] back to full [0..255] range. Threshold values were chosen manually for Hoechst and marker channels for each image in order to remove background from the images. For all markers except HIF1α, total protein expression levels in a tissue were quantified as a relative number of cells expressing the protein, i.e.

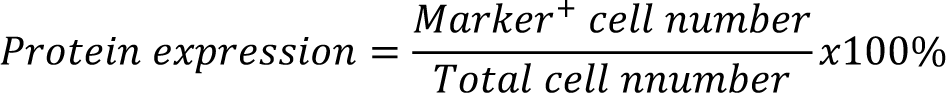

On the assumption that cell size is constant across the samples, the relative number of cells was estimated as a ratio of areas occupied by cells on marker and Hoechst channels. Corresponding areas were calculated by counting all non-zero pixels on marker and Hoechst channels. For statistical comparison, results for each marker were pooled from all experiments.

As HIF1α marker is constitutively expressed in cells under normoxic conditions, its expression level in a tissue was estimated as the average fluorescence intensity on the marker channel. Threshold values were kept constant for organoids, non-perfused and perfused samples obtained in the same experiment; expression values obtained in the experiment were normalized to organoid value and normalized data from different experiments were averaged.

All results are expressed as mean ± SEM. Statistical comparisons between two data groups were analyzed using unpaired two-tailed Student’s *t*-test with a 95% confidence interval (Origin Pro v9.5, OriginLab Inc.). Significance was marked on plots by *, ** and *** for p<0.05, *p* < 0.01, and *p* < 0.001 correspondingly. If not stated otherwise, the statistical significance was assessed for comparisons of perfused and non-perfused samples to organoid control samples and denoted by “p_Control_”.

### Organoid dissociation and single-cell RNA sequencing

For dissociation, the microfluidic grids of non-perfused and perfused samples were removed from the perfusion chips and, along with the control organoids, placed temporarily in DMEM/F12 media in independent 15 mL Falcon tubes until all 3 samples were ready for the next step (time < 5 min). The DMEM was then replaced with 1 mL of pre-warmed TrypLE Express (12605010, Gibco) at 37°C. The samples were transferred to a warm water bath at 37°C for 7.5 minutes with gentle agitation every 1 minute after the first 3 minutes. A visual inspection was performed using an inverted microscope to ensure complete dissociation of tissues and the presence of single cells. The 1 mL solution was introduced to 9 mL of DMEM supplemented with 20% FBS for TrypLE Express neutralization, and centrifuged at 500 rcf for 5 minutes. The pellet was resuspended in 200 uL of N2B27 media and put on ice. This dissociation protocol yielded an average viability of above 80% across all samples.

Library preparations for the scRNAseq was performed using 10X Genomics Chromium Single Cell 3’ Kit, v3 (10X Genomics, Pleasanton, CA, USA). The cell count and the viability of the samples were accessed using LUNA dual fluorescence cell counter (Logos Biosystems) and a targeted cell recovery of 6000 cells was aimed for all the samples. Post cell count and QC, the samples were immediately loaded onto the Chromium Controller. Single cell RNAseq libraries were prepared using manufacturers recommendations (Single cell 3’ reagent kits v3 user guide; CG00052 Rev B), and at the different check points, the library quality was accessed using Qubit (ThermoFisher) and Bioanalyzer (Agilent). Single cell libraries were sequenced using paired-end sequencing workflow and with recommended 10X; v3 read parameters (28-8-0-91 cycles). The data generated from sequencing was de-multiplexed and mapped against human genome reference using CellRanger v3.0.2.

### Single Cell RNA sequencing Data Processing

We have sequenced 4,818, 5,453, and 2,804 cells for non-perfused, perfused and control organoids samples respectively for a total of 13,075 cells, which were reduced after QC steps to 8,625 cells with an average of 1,852 detected genes per cell. Data processing and subsequent steps were performed using the Seurat^72^ tool for single cell genomics version 3 in R version 4.0.3. A filtering step was performed to ensure the quality of the data, where the counts of mitochondrial reads and total genes reads were assessed. Cells with more than 15% of identifiable genes rising from the mitochondrial genome were filtered out. Similarly, cells having fewer identifiable genes than 200 (low quality) and above 7,500 (probable doublets) were filtered out. Data normalization was performed, followed by the identification of 2,000 highly variable genes using the FindVariableFeatures(). S-phase and G2M-phase cell cycle regression was performed to allow cell clustering purely on cell identity and fate, which otherwise was biased by cell cycle phases. Auto scaling of the data was performed and described using principal component analysis (PCA) using the RunPCA() function.

### Correlation analysis

Correlation heatmap between samples was performed in R using the top 100 marker genes for each sample using the FindAllMarkers() function followed by the selection of the top 100 marker genes using the top_n() function with n = 100 and wt = avg.logFC. The correlation analysis was generated using the cor() function using the Pearson method and the heatmap using the heatmap.2() function in R.

### Data Clustering

Graph-based clustering using the FindNeighbors() function (using top 15 principal components (PCs)) and FindCluster() function (resolution = 0.5) was performed to group cells based on their transcriptional profiles. No batch correction was performed on the data set. Data visualization was performed using the Uniform Manifold Approximation and Projection (UMAP) dimensionality reduction technique using the RunUMAP package while employing the top 15 PCs identified in the previous PCA step. Cluster annotation was based on hallmark genes to identify Neuroepithelial cluster (PAX6, PAX7, PAX3), Pluripotent-Neuroepithelial transitioning (P-NE) cells (WNT4, IRX1, IRX2), Proliferating cells (CDC6, PTN, MKI67, TOP2A), medium mitochondrial gene content and low glycolysis pluripotent cells (MKI67, TOP2A, with the expression of NANOG, POU5F1, DPPA4), Pluripotent cells (NANOG, POU5F1, DPPA4), highly glycolytic pluripotent cells that link to the Cycling pluripotent and Pluripotent cells but include the expressions (LDHA, ENO1, HK2), a cluster undergoing EMT/migration with hypoxic markers (EPCAM, VIM, TWIST2, VEGFA) and finally a stressed cluster (FOS, HIF1A, JUN). One cluster was manually removed due to very low unique molecular identifier (UMI) count, after which the data was reprocessed to account for the removal of the cluster.

### Pseudotime trajectories

Trajectory analysis was performed using Monocle 3^73^ on R by employing the monocole3 and SeuratWrappers libraries. The Seurat obtained object containing all combined samples was passed to the Monocle 3 pipeline using the as.cell_data_set() function followed by processing using the cluster_cells() function. The pseudotime trajectory was inferred by first using the subset function on the only partition detected, followed by the learn_graph() function. Cells were colored in their inferred pseudotime using the which.max() function with the FetchData() (AVP) function, followed by the order_cells() function with the root_cell = max.avp. The UMAP superimposed with inferred pseudotime trajectory was generated using the plot_cells() function.

### Gene-set analysis and hierarchical clustering

Gene-sets were created using the AddModuleScore() function in R. three gene-set were created to explore the dataset, pluripotency gene-set (*POU5F1, NANOG, CDH1*), glycolysis gene-set (*ALDOA, BPGM, ENO1, ENO2, GPI, HK1, HK2, PFKL, PFKM, PGAM1, PGK1, PKM, TP11*), and a neural progenitor gene-set (*PAX3, PAX6, PAX7, OTX2, CDH2*). The % mitochondrial genes were obtain as evaluated by the Seurat pipeline. Each cell was evaluated and scored and its expression plotted on UMAPS. Cluster level hierarchical clustering of the gene-set scores and % mitochondrial genes was employed using R package heatmap.2 after scaling the expression across rows (clusters).

### Design and fabrication of soft micro-fluidic grids

The microfluidic grids were designed as a multitude of parallel capillaries stemming from a common reservoir. The dimensions of the grid (2.6×2.6×1.5mm) were chosen as a compromise between size and the fabrication time. The grids were micro-fabricated on custom baseplates using high-resolution 3D printer (Photonic Professional GT2, Nanoscribe GmbH) equipped with 2-photon femtosecond laser. CAD model of the capillary grid was pre-processed by DeScribe software (Nanoscribe GmbH) to produce a printing job with defined printing parameters. A custom-formulation photopolymerizable resin was used to print microfluidic grids which contained 2% 2-Benzyl-2-(dimethylamino)-4’-morpholinobutyrophenone (Irgacure 369), 7% propylene glycol methyl ether acetate (PGMEA), 36% poly(ethylene glycol) diacrylate, MW700 (PEGDA 700), 25% pentaerythritol triacrylate (PETA), 20% Triton X-100 and 10% water. Irgacure 369 was purchased from Tokyo Chemical Industry Ltd, the other components were from Sigma-Aldrich.

The microfluidic grids were printed on custom plates 3D printed on Formlabs Form 2 printer from a biocompatible resin (Dental SG, Formlabs Inc), post-cured by UV light (Formcure station, Formlabs Inc) for 2h at 80°C and washed in 2-propanol for 3-4 days with daily change of the solvent. The plates had 1mm thickness and 10mm diameter, with 0.5mm perfusion holes. The grid printing process was set up such that the inlets in the microfluidic grid were aligned with the perfusion holes in a baseplate. After fabrication, baseplate-grid assemblies were washed during 2 days in PGMEA, 12 days in 2-propanol and then kept in PBS until use. The solvents were refreshed every 1-2 days and PBS was refreshed daily for the first week and every 2-3 days afterwards.

### Fabrication of perfusion chips and setting up perfusion

The perfusion chips were implemented as an assembly as shown in Supplementary Fig. 1. Two microfluidic grids were sandwiched between two polydimethylsiloxane (PDMS) custom-profiled blocks. Two metal plates clamped with screws provided a tight junction between PDMS blocks and cover slips, thereby forming fluidic channels within the chip. The chips were designed to only allow contact of the medium with PDMS blocks and cover slips, thereby isolating the flow from contact with other materials of the chip. Guiding frames were used to simplify the alignment of the PDMS blocks during the chip assembly process.

The PDMS blocks were made by casting liquid PDMS compound into custom-designed plastic molds and curing overnight at 80°C. The plastic molds were fabricated by stereo-lithography (SLA) 3D printer Form 2 (Formlabs Inc.) using a bio-compatible Formlabs Dental SG resin. The printed molds were post-processed by thorough washing in 2-propanol and post-cured for 90 min at 80°C (Formcure, Formlabs Inc.). Metal plates were made from 1mm stainless steel sheet by laser cutting and stainless steel screws were used for clamping the assembly. The guiding frames were 3D printed and cover slips were glued to the metal plates with a bio-compatible UV/heat curable epoxy (NOA 86H, Norland) to provide a final two-part setup shown in Supplementary Fig. 1d.

Two chambers of the chip, containing perfused and non-perfused grids, a peristaltic pump and a medium reservoir (50ml Falcon tube) were connected by flexible PVC tubing (ID/OD 0.8/1.6mm) in series (Supplementary Fig. 1e). The medium was taken from the reservoir by the pump and fed through chambers containing the non-perfused and then the perfused grid, after which the medium was sent back to the reservoir for CO_2_/O_2_ equilibration. A 0.22um filter was connected between the two chambers of the chip to protect capillaries of the perfused grid from potential clogging with tissue debris. The medium reservoir and the chip were kept in a CO2 incubator, while the peristaltic pump remained outside to protect electronics from humidity and to ensure proper cooling. The medium reservoir and the chip were installed on a custom made 3D printed holder (Supplementary Fig. 1f) to simplify transitions between a CO2 incubator and a laminar hood. At day 0 of an experiment, all parts of the perfusion were sterilized by ethanol and UV light in a laminar hood, connected together and filled with equilibrated medium. The microfluidic grids, seeded with hPSC spheroids, were installed in the chip (Supplementary Fig. 1d) and the chip was clamped by screws (Supplementary Fig.1f-i). Air bubbles were removed from the system and the perfusion started at approx. 400µL/min flow rate (Supplementary Fig. 1,f-iii).

Multi-grid perfusion chips were designed to provide an equal multiplexed perfusion for all microfluidic grids. The fluidic inlet of the chip, via three steps of bifurcations, was split into eight fluidic channels feeding eight wells. All fluidic channels had equal length and cross-section and, therefore, equal hydraulic resistance. The multi-grid chip were fabricated by stereo-lithography (SLA) 3D printer Form 2 (Formlabs Inc.) using Formlabs Dental SG resin. The printed chips were post-processed by thorough washing in 2-propanol and post-cured for 90 min at 60°C. 7-8mm pieces of syringe needles (0.9mm) were cut by hand drill/cutter (Precision drill, Proxxon Inc.) and glued into chip inlets by a biocompatible UV/heat curable epoxy glue to serve as connectors for flexible tubing. This assembly was again post-cured in FormCure station for 120 min at 80°C to achieve complete hardening of the glue. To integrate microfluidic grids, a custom profile gaskets, cut from 0.5mm PDMS sheet by a laser cutter, were placed under the grid substrates to provide a liquid-tight contact and the substrates were fixed in place by 3D printed fasteners.

## Data availability

All raw sequencing data, and the combined processed and metadata files generated in this study are available at GEO. The accession number for the reported data is GSE181290. This study did not generate any unique code. For review purposes please use the following token (ybchyyoqfpejzgv) to access the sequencing data at: https://www.ncbi.nlm.nih.gov/geo/query/acc.cgi?acc=GSE181290. The ImageJ Java code used for analysis of the image data are available upon request.

## Supplementary Figures

**Supplementary Figure 1.**
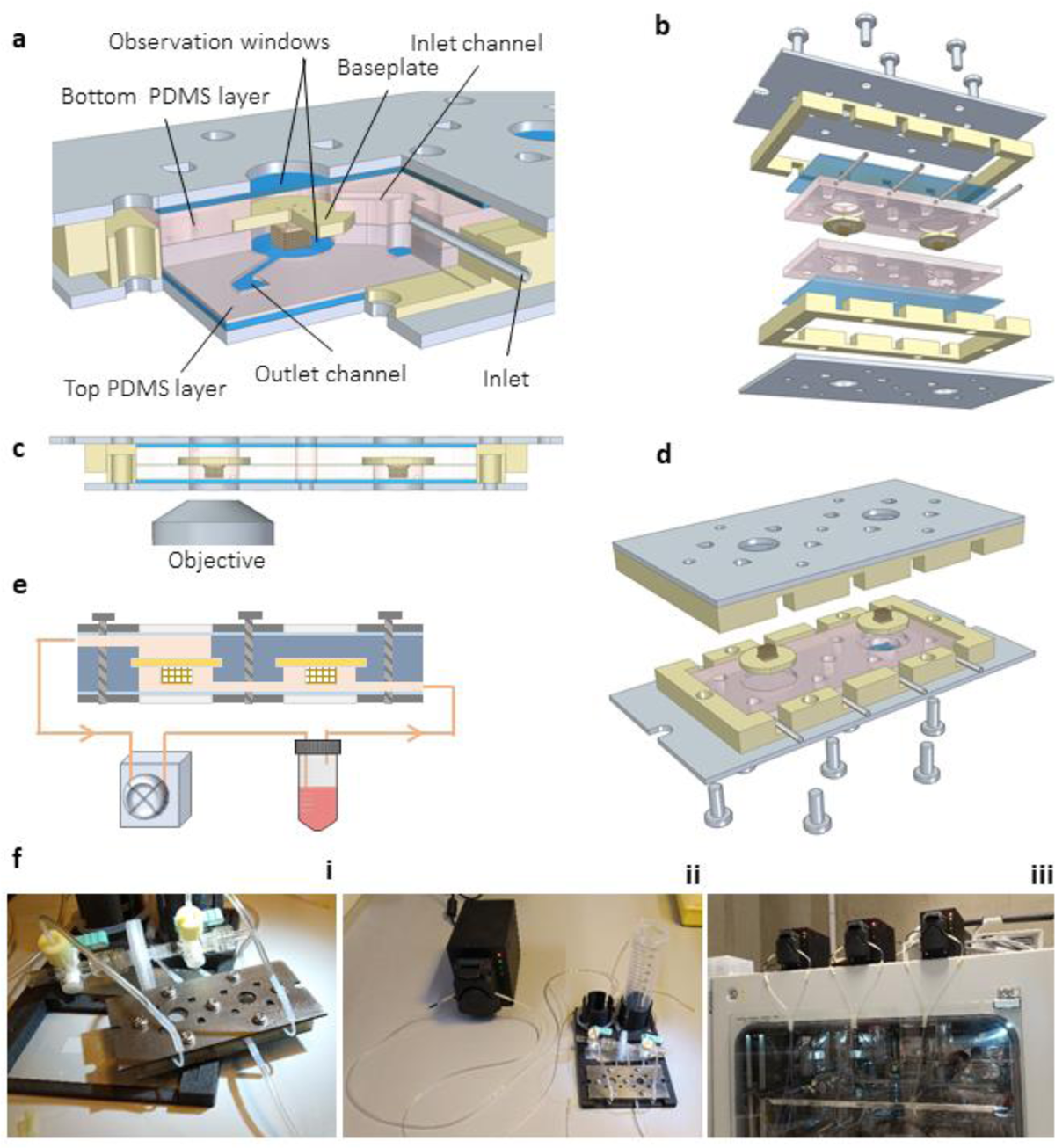
Assembly and operation of the chip device. (**a**) Cutaway view of the fully assembled culture chip, the chamber with perfused capillary grid is shown. Cell culture medium enters the culture chamber through the inlet (1mm syringe needle, cut into 15mm pieces), passes through the capillary grid and leaves the chamber through outlet (not shown). (**b**) Exploded view of the chip. (**c**) Side view cross section of the chip. Tissue constructs can be observed through the glass windows with inverted or upright microscopes. The view also demonstrates that the cell culture medium only comes into contact with PDMS(pink) and glass(blue) parts of the chip. (**d**) During the assembly, a user needs to handle only two pre-assembled “halves” of the chip. (**e**) Schematic representation of the perfusion system driven by a peristaltic pump, with culture medium circulating between the perfusion chip and the medium reservoir. Here, the grid in the right chamber is surrounded by a constant medium flow but the flow through its capillaries is absent, while in the downstream grid (left chamber) the flow passes through the capillaries and diffuses across the walls into the tissue. (**f**) Photographs of fully assembled chip and perfusion system. The non-perfused and perfused chambers are connected by external tubing (left), with 0.22µm filter connected inline between the chambers. The chip and the medium reservoir resides in a custom 3D-printed holder (middle) and connected to a peristaltic pump installed outside of CO_2_ incubator (right). The image shows three experiments running in parallel.

**Supplementary Figure 2.**
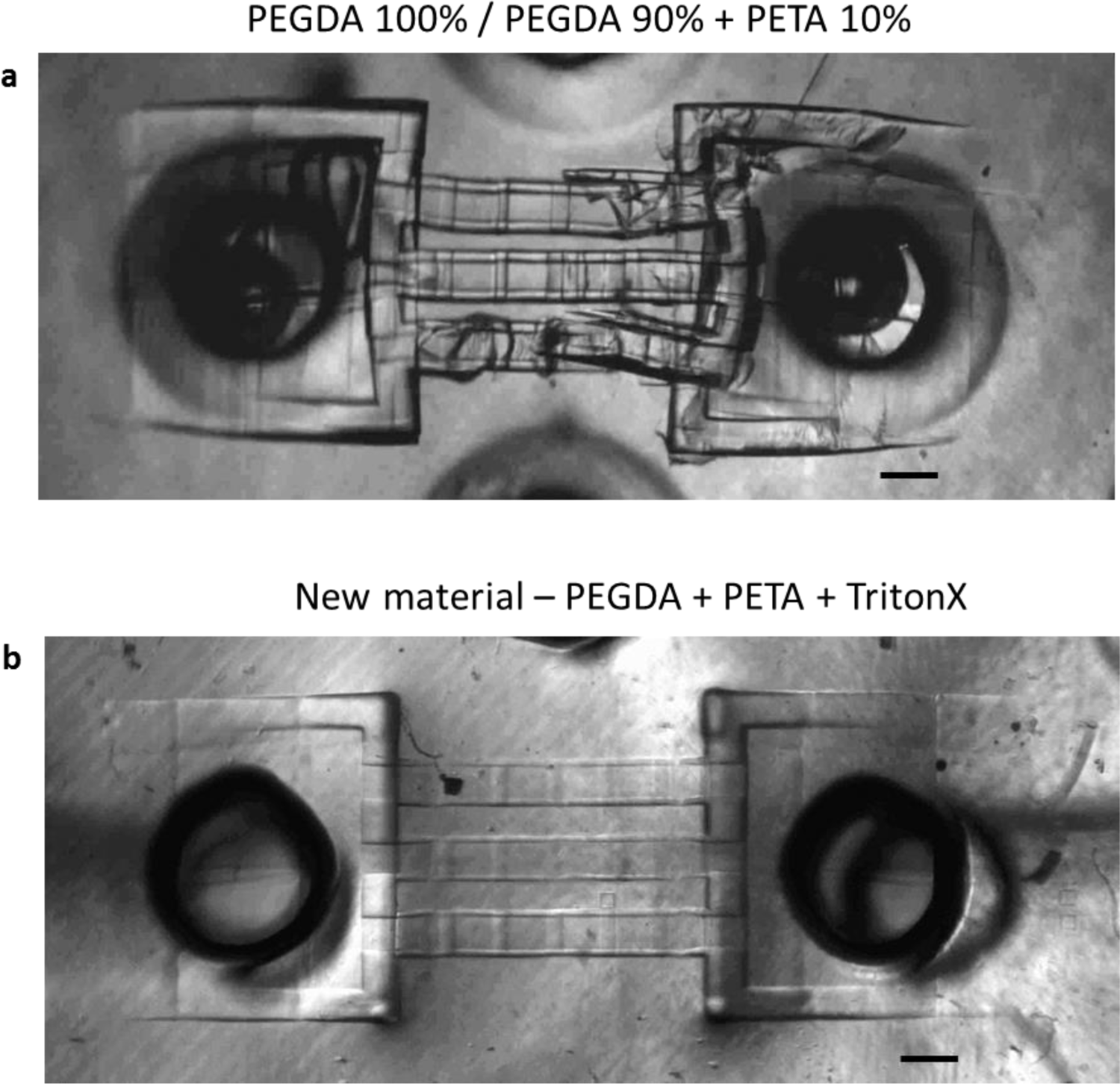
Optimization of printable material properties. (**a**) Test structure printed from PEGDA or PEGDA 90%/PETA 10% mixture. Both ends of the part are connected to a baseplate. When submerged to cell culture medium, swelling of the material leads to a breakage of the part. (**b**) Our custom formulation resin effectively prevents swelling in aqueous media and preserves the geometry of the micro-fabricated structures. Scale bar 100µm.

**Supplementary Figure 3.**
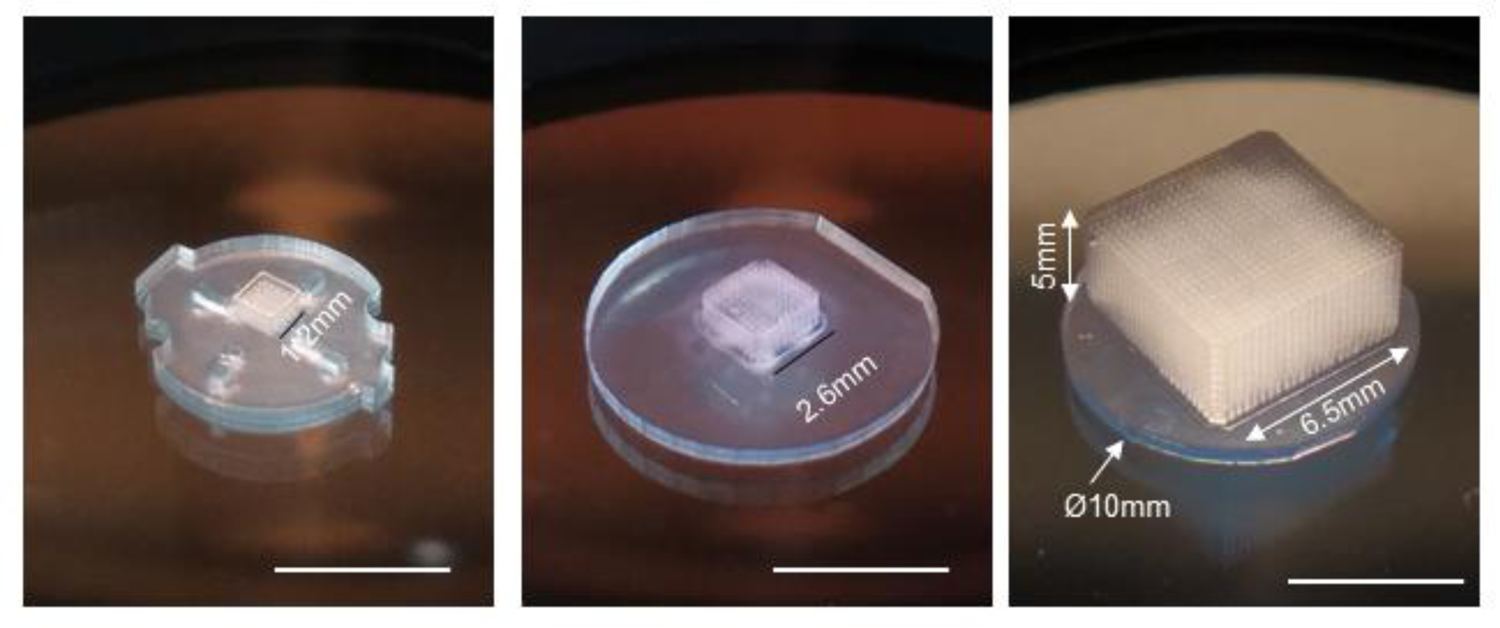
Two-photon stereo-lithography enables precision fabrication in a wide range of scales. Microfluidic grids of different dimensions (in mm): 1.2mm x 1.2mm x 1.2mm (left), 2.6mm x 2.6 mm x 1.5mm (middle), 6.5mm x 6.5mm x 5mm (right). Perfusion vessels in each grid have identical 50µm diameter and inter-vessel distance of 250µm. Scale bar 5 mm.

**Supplementary Figure 4.**
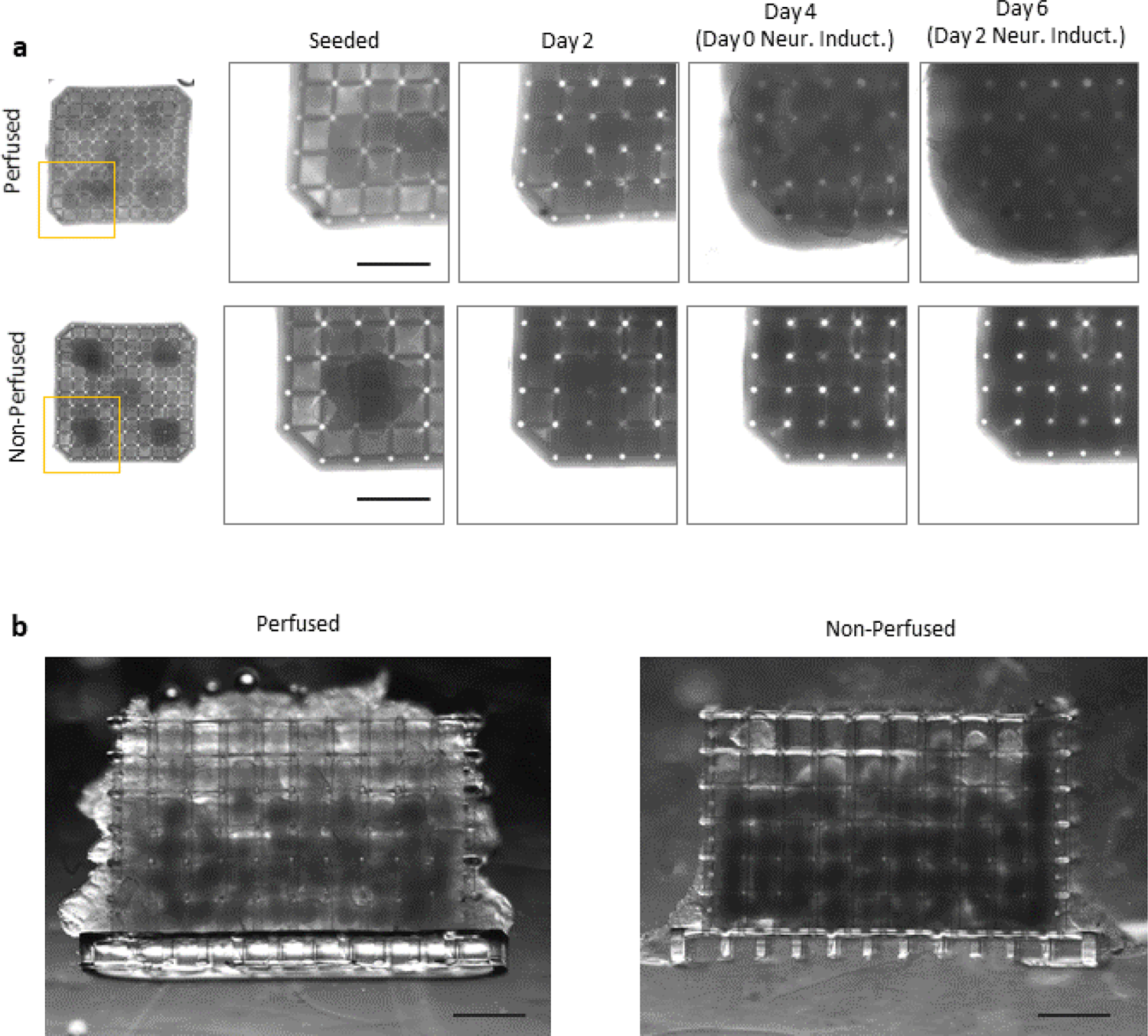
Daily brightfield imaging of neural tissue in perfused and non-perfused chips. (**a**) Bright field images of the perfused (top) and non-perfused (bottom) tissue constructs taken every 2 days during the culturing protocol. (**b**) Live sections of perfused (left) and non-perfused (right) tissue constructs taken at the end of the culturing protocol. Scale bar 500µm.

**Supplementary Figure 5.**
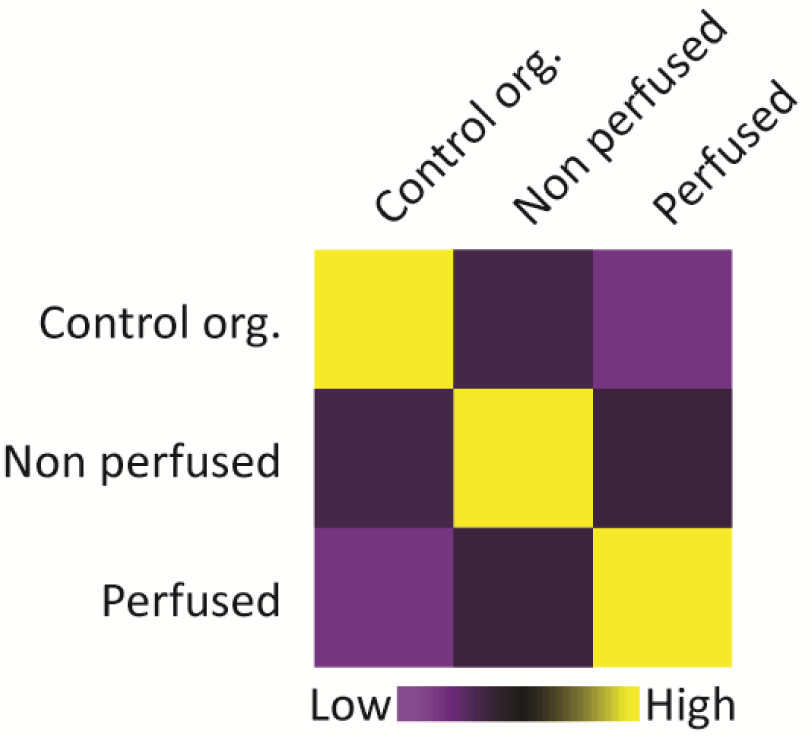
Correlation analysis between perfused-, non-perfused neural tissue constructs and conventional neural organoid culture. Correlation analysis using the top 100 differentially expressed marker genes for each of the three experimental conditions.

**Supplementary Figure 6.**
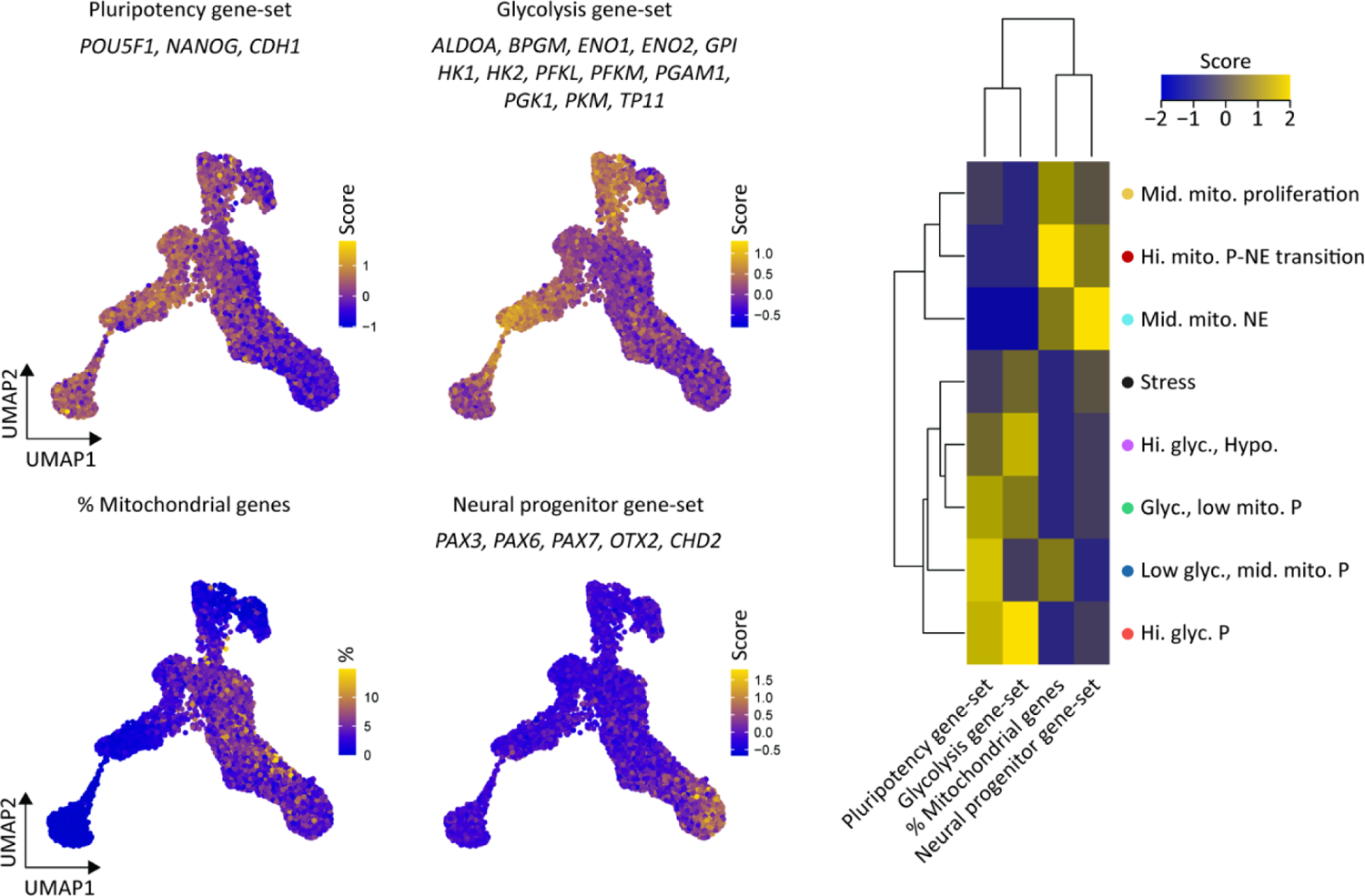
Gene-set analysis for various processes. Combined dataset UMAP with scores for pluripotent, glycolysis and neural progenitor gene-set as well as the % mitochondrial genes. Hierarchical clustering of gene-set scores and % mitochondrial genes. Score is column scaled.

**Supplementary Figure 7.**
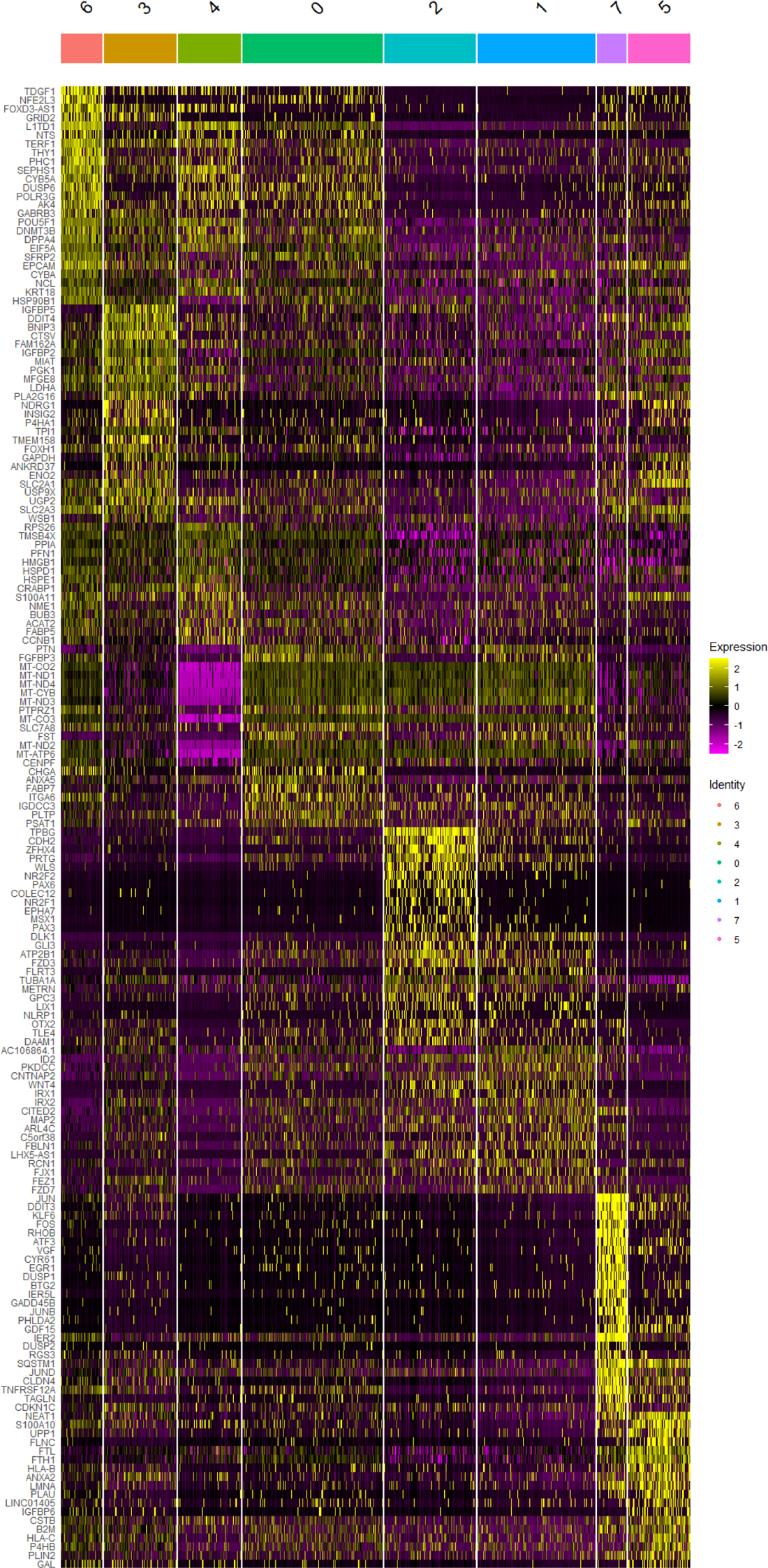
Gene expression for unannotated hNTO clusters with the top 25 marker genes for each cluster.

**Supplementary Figure 8.**
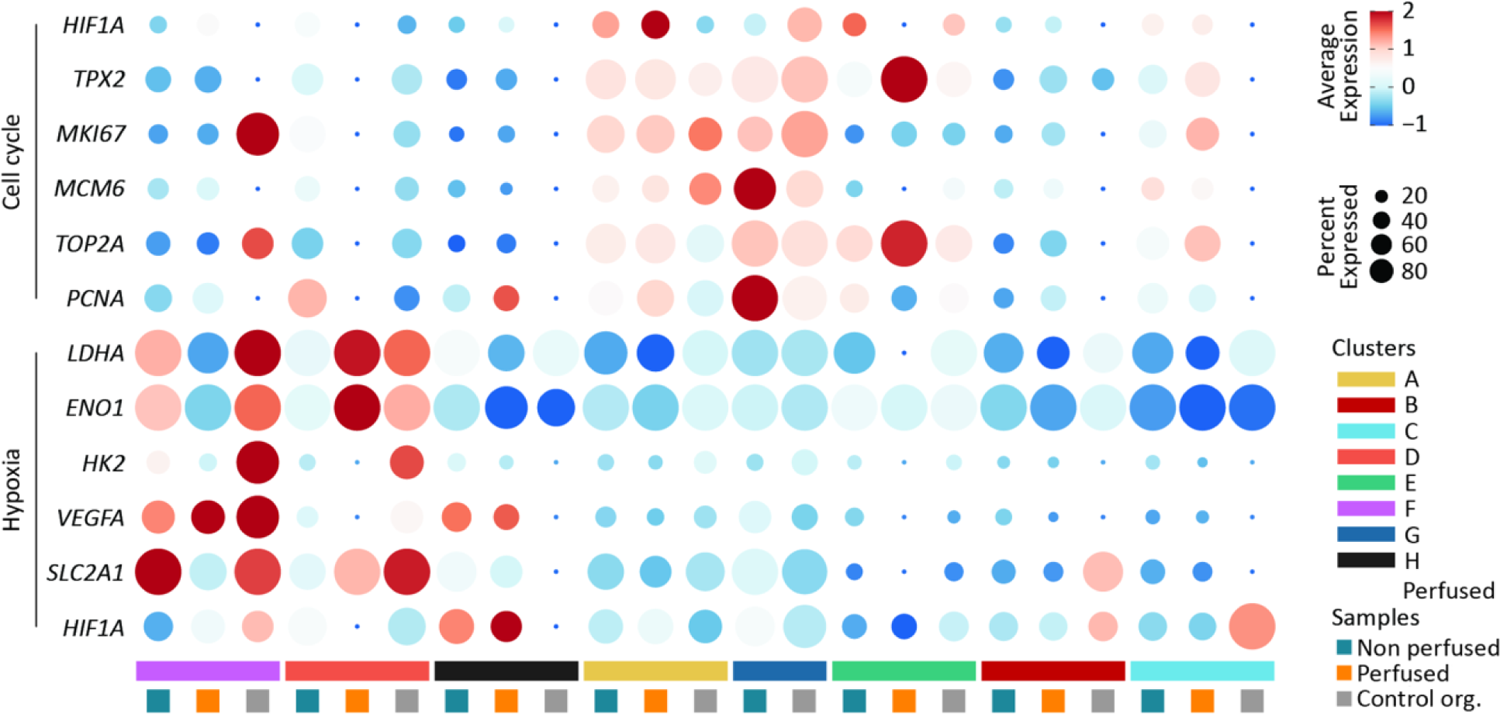
Cluster-specific analysis of scRNAseq dataset. Dot-plot heatmap of hypoxia and cell cycle markers for each identified cluster. The average gene expression is represented by the color intensity of each dot, whereas the dot size represents the percentage of the gene-expressing cells for each sample within each cluster.

**Supplementary Figure 9.**
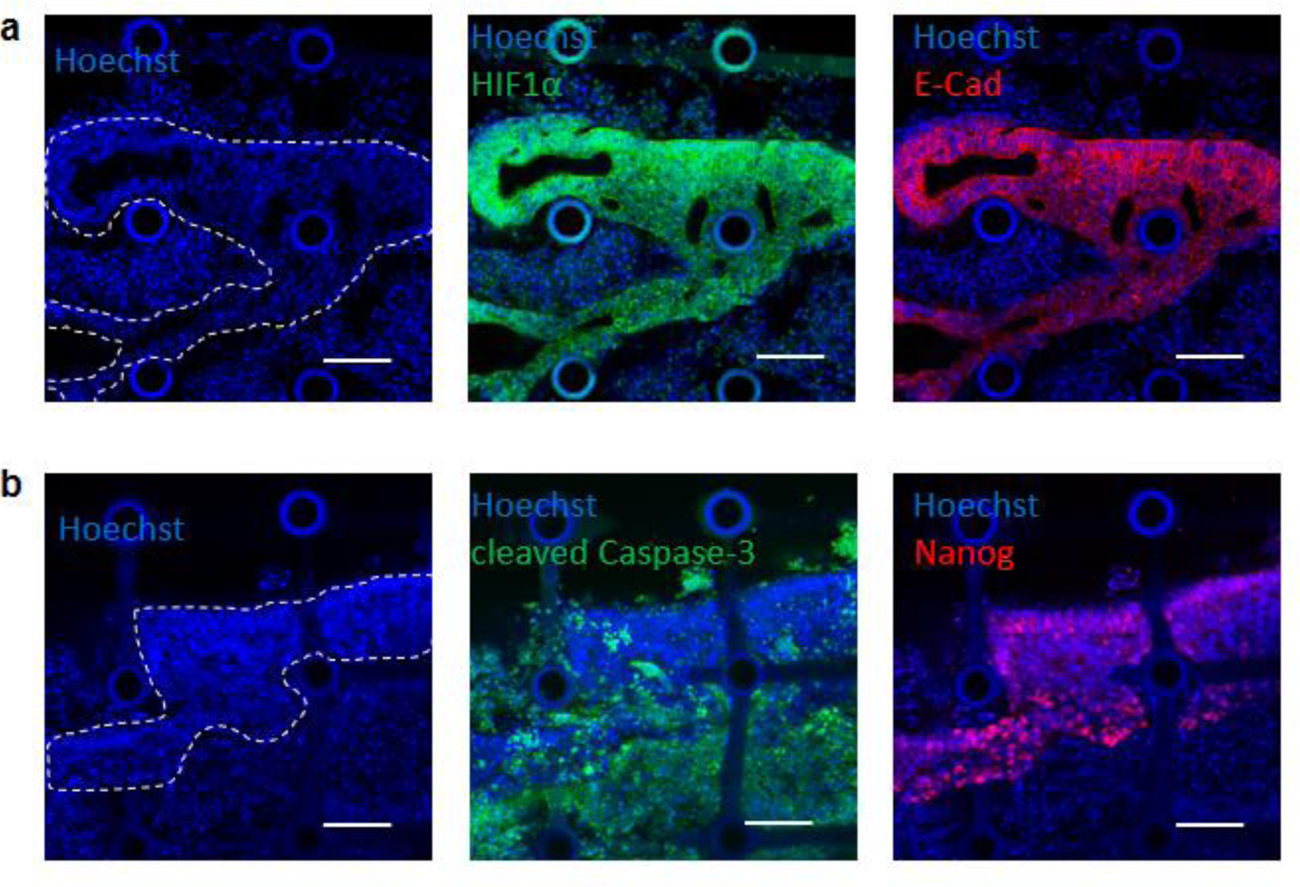
Difference in localized expression of hypoxia marker HIF1α and apoptosis marker cleaved Caspase-3. (**a**) HIF1α expression is localized to regions containing intact cell bodies (middle), evidenced by intact nuclei in the outlined region (left) as well as well-defined cytoplasmic regions stained with E-Cad antibody (right). (**b**) Cleaved Caspase-3 expression (middle) is localized to regions with apoptotic cell bodies evidenced by Hoechst stain of fragmented nuclei (outside of the outlined live region on the left image). Scale bar 100µm.

**Supplementary Figure 10.**
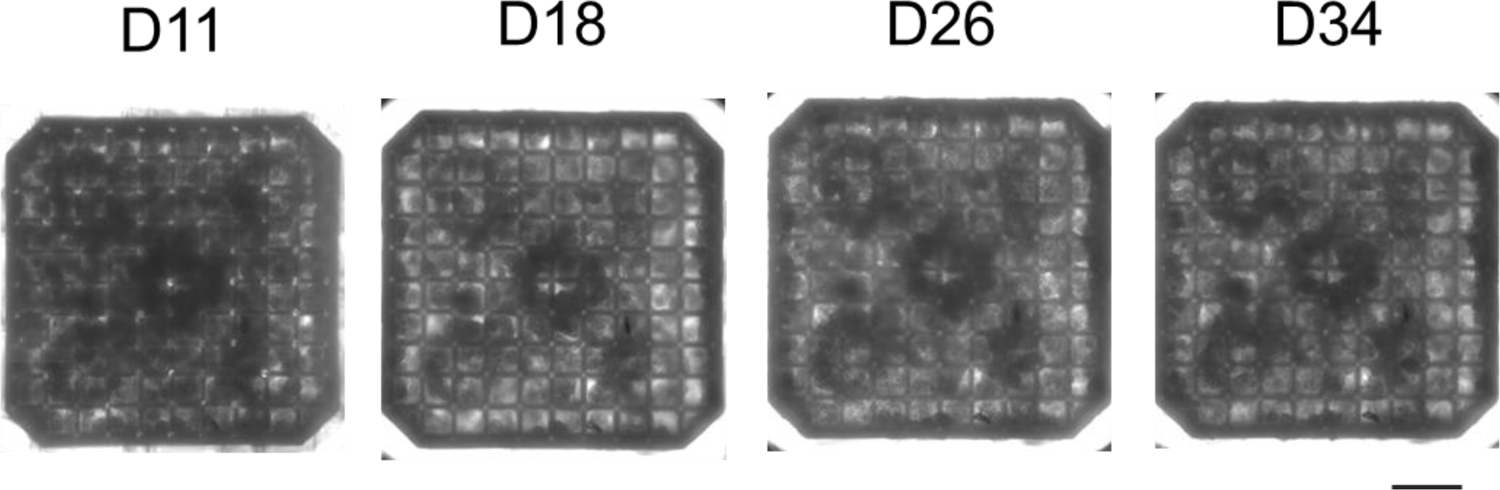
Brightfield images of perfused liver cultures. Bright field images of the perfused liver-like tissue constructs taken during tissue culturing. Scale bar 500µm.

## References

1. Kaushik, G., Ponnusamy, M. P. & Batra, S. K. Concise Review: Current Status of Three-Dimensional Organoids as Preclinical Models: 3D Organoid Culture as a Tool for Research. Stem Cells 36, 1329–1340 (2018).

2. Ollé-Vila, A., Duran-Nebreda, S., Conde-Pueyo, N., Montañez, R. & Solé, R. A morphospace for synthetic organs and organoids: the possible and the actual. Integr. Biol. 8, 485–503 (2016).

3. Van Norman, G. A. Limitations of Animal Studies for Predicting Toxicity in Clinical Trials. JACC: Basic to Translational Science 4, 845–854 (2019).

4. Gilbert, S. F. Developmental biology. (Sinauer Associates, 2000).

5. Grebenyuk, S. & Ranga, A. Engineering Organoid Vascularization. Front. Bioeng. Biotechnol. 7, 39 (2019).

6. Nashimoto, Y. et al. Integrating perfusable vascular networks with a three-dimensional tissue in a microfluidic device. Integrative Biology 9, 506–518 (2017).

7. Salmon, I. et al. Engineering neurovascular organoids with 3D printed microfluidic chips. http://biorxiv.org/lookup/doi/10.1101/2021.01.09.425975 (2021) doi:10.1101/2021.01.09.425975.

8. Rajasekar, S. et al. IFlowPlate—A Customized 384-Well Plate for the Culture of Perfusable Vascularized Colon Organoids. Adv. Mater. 32, 2002974 (2020).

9. Sugihara, K. et al. A new perfusion culture method with a self-organized capillary network. PLoS ONE 15, e0240552 (2020).

10. Takebe, T. et al. Vascularized and functional human liver from an iPSC-derived organ bud transplant. Nature 499, 481–484 (2013).

11. Mansour, A. A. et al. An in vivo model of functional and vascularized human brain organoids. Nature Biotechnology 36, 432–441 (2018).

12. Zhu, W. et al. 3D printing of functional biomaterials for tissue engineering. Current Opinion in Biotechnology 40, 103–112 (2016).

13. Xie, M. et al. Electro-Assisted Bioprinting of Low-Concentration GelMA Microdroplets. Small 15, 1804216 (2019).

14. Cui, X., Boland, T., D.D’Lima, D. & K. Lotz, M. Thermal Inkjet Printing in Tissue Engineering and Regenerative Medicine. Recent Patents on Drug Delivery & Formulation 6, 149–155 (2012).

15. Dababneh, A. B. & Ozbolat, I. T. Bioprinting Technology: A Current State-of-the-Art Review. Journal of Manufacturing Science and Engineering 136, 061016 (2014).

16. Gudapati, H., Dey, M. & Ozbolat, I. A comprehensive review on droplet-based bioprinting: Past, present and future. Biomaterials 102, 20–42 (2016).

17. Christensen, K. et al. Freeform inkjet printing of cellular structures with bifurcations: Approach Freeform Fabrication of Bifurcated Cellular Structures by Using a Liquid Support-Based Inkjet Printing Approach. Biotechnology and Bioengineering 112, 1047–1055 (2015).

18. Nakamura, M. et al. Ink Jet Three-Dimensional Digital Fabrication for Biological Tissue Manufacturing: Analysis of Alginate Microgel Beads Produced by Ink Jet Droplets for Three Dimensional Tissue Fabrication. Journal of Imaging Science and Technology 52, 060201 (2008).

19. Zhu, W. et al. Direct 3D bioprinting of prevascularized tissue constructs with complex microarchitecture. Biomaterials 124, 106–115 (2017).

20. Ma, X. et al. Deterministically patterned biomimetic human iPSC-derived hepatic model via rapid 3D bioprinting. Proceedings of the National Academy of Sciences 113, 2206–2211 (2016).

21. Huang, T. Q., Qu, X., Liu, J. & Chen, S. 3D printing of biomimetic microstructures for cancer cell migration. Biomedical Microdevices 16, 127–132 (2014).

22. Singh, N. K. et al. Three-dimensional cell-printing of advanced renal tubular tissue analogue. Biomaterials 232, 119734 (2020).

23. Gao, Q. et al. 3D printing of complex GelMA-based scaffolds with nanoclay. Biofabrication 11, 035006 (2019).

24. Jia, W. et al. Direct 3D bioprinting of perfusable vascular constructs using a blend bioink. Biomaterials 106, 58–68 (2016).

25. Xu, C., Chai, W., Huang, Y. & Markwald, R. R. Scaffold-free inkjet printing of three-dimensional zigzag cellular tubes. Biotechnology and Bioengineering 109, 3152–3160 (2012).

26. Kinoshita, K., Iwase, M., Yamada, M., Yajima, Y. & Seki, M. Fabrication of multilayered vascular tissues using microfluidic agarose hydrogel platforms. Biotechnology Journal (2016) doi:10.1002/biot.201600083.

27. Roudsari, L. C., Jeffs, S. E., Witt, A. S., Gill, B. J. & West, J. L. A 3D Poly(ethylene glycol)-based Tumor Angiogenesis Model to Study the Influence of Vascular Cells on Lung Tumor Cell Behavior. Scientific Reports 6, 32726 (2016).

28. Zhang, Y. S. et al. Bioprinting 3D microfibrous scaffolds for engineering endothelialized myocardium and heart-on-a-chip. Biomaterials 110, 45–59 (2016).

29. Meyer, W. et al. Soft Polymers for Building up Small and Smallest Blood Supplying Systems by Stereolithography. Journal of Functional Biomaterials 3, 257–268 (2012).

30. Huber, B. et al. Blood-Vessel Mimicking Structures by Stereolithographic Fabrication of Small Porous Tubes Using Cytocompatible Polyacrylate Elastomers, Biofunctionalization and Endothelialization. Journal of Functional Biomaterials 7, 11 (2016).

31. Kang, H.-W. et al. A 3D bioprinting system to produce human-scale tissue constructs with structural integrity. Nature Biotechnology 34, 312–319 (2016).

32. Compaan, A. M., Song, K., Chai, W. & Huang, Y. Cross-Linkable Microgel Composite Matrix Bath for Embedded Bioprinting of Perfusable Tissue Constructs and Sculpting of Solid Objects. ACS Appl. Mater. Interfaces 12, 7855–7868 (2020).

33. Miller, J. S. et al. Rapid casting of patterned vascular networks for perfusable engineered three-dimensional tissues. Nature Materials 11, 768–774 (2012).

34. Kolesky, D. B., Homan, K. A., Skylar-Scott, M. A. & Lewis, J. A. Three-dimensional bioprinting of thick vascularized tissues. Proceedings of the National Academy of Sciences 113, 3179–3184 (2016).

35. Wu, W., DeConinck, A. & Lewis, J. A. Omnidirectional Printing of 3D Microvascular Networks. Advanced Materials 23, H178–H183 (2011).

36. Bertassoni, L. E. et al. Hydrogel bioprinted microchannel networks for vascularization of tissue engineering constructs. Lab Chip 14, 2202–2211 (2014).

37. Subbiah, R. et al. Prevascularized hydrogels with mature vascular networks promote the regeneration of critical-size calvarial bone defects in vivo A short running title: Prevascularized hydrogels repair bone defects. J Tissue Eng Regen Med (2021) doi:10.1002/term.3166.

38. Skylar-Scott, M. A. et al. Biomanufacturing of organ-specific tissues with high cellular density and embedded vascular channels. Sci. Adv. 5, eaaw2459 (2019).

39. Silvestri, V. L. et al. A Tissue-Engineered 3D Microvessel Model Reveals the Dynamics of Mosaic Vessel Formation in Breast Cancer. Cancer Res 80, 4288–4301 (2020).

40. Applegate, M. B. et al. Laser-based three-dimensional multiscale micropatterning of biocompatible hydrogels for customized tissue engineering scaffolds. Proceedings of the National Academy of Sciences 112, 12052–12057 (2015).

41. Oujja, M. et al. Three dimensional microstructuring of biopolymers by femtosecond laser irradiation. Applied Physics Letters 95, 263703 (2009).

42. Sarig-Nadir, O., Livnat, N., Zajdman, R., Shoham, S. & Seliktar, D. Laser Photoablation of Guidance Microchannels into Hydrogels Directs Cell Growth in Three Dimensions. Biophysical Journal 96, 4743–4752 (2009).

43. Brandenberg, N. & Lutolf, M. P. In Situ Patterning of Microfluidic Networks in 3D Cell-Laden Hydrogels. Advanced Materials 28, 7450–7456 (2016).

44. Skylar-Scott, M. A., Liu, M.-C., Wu, Y. & Yanik, M. F. Multi-photon microfabrication of three-dimensional capillary-scale vascular networks. in (eds. von Freymann, G., Schoenfeld, W. V. & Rumpf, R. C.) 101150L (2017). doi:10.1117/12.2253520.

45. Kloxin, A. M., Kasko, A. M., Salinas, C. N. & Anseth, K. S. Photodegradable Hydrogels for Dynamic Tuning of Physical and Chemical Properties. Science 324, 59–63 (2009).

46. Kloxin, A. M., Tibbitt, M. W., Kasko, A. M., Fairbairn, J. A. & Anseth, K. S. Tunable Hydrogels for External Manipulation of Cellular Microenvironments through Controlled Photodegradation. Advanced Materials 22, 61–66 (2010).

47. Tibbitt, M. W., Kloxin, A. M., Dyamenahalli, K. U. & Anseth, K. S. Controlled two-photon photodegradation of PEG hydrogels to study and manipulate subcellular interactions on soft materials. Soft Matter 6, 5100 (2010).

48. Kim, J., Kong, J. S., Han, W., Kim, B. S. & Cho, D.-W. 3D Cell Printing of Tissue/Organ-Mimicking Constructs for Therapeutic and Drug Testing Applications. IJMS 21, 7757 (2020).

49. Ahadian, S. et al. Organ-On-A-Chip Platforms: A Convergence of Advanced Materials, Cells, and Microscale Technologies. Advanced Healthcare Materials 1700506 (2017) doi:10.1002/adhm.201700506.

50. Mittal, R. et al. Organ-on-chip models: Implications in drug discovery and clinical applications. J Cell Physiol 234, 8352–8380 (2019).

51. Van Norman, G. A. Limitations of Animal Studies for Predicting Toxicity in Clinical Trials. JACC: Basic to Translational Science 5, 387–397 (2020).

52. Gaetani, R. et al. Epicardial application of cardiac progenitor cells in a 3D-printed gelatin/hyaluronic acid patch preserves cardiac function after myocardial infarction. Biomaterials 61, 339–348 (2015).

53. Norona, L. M., Nguyen, D. G., Gerber, D. A., Presnell, S. C. & LeCluyse, E. L. Editor’s Highlight: Modeling Compound-Induced Fibrogenesis In Vitro Using Three-Dimensional Bioprinted Human Liver Tissues. Toxicol. Sci. 154, 354–367 (2016).

54. Klein, F. et al. Two-Component Polymer Scaffolds for Controlled Three-Dimensional Cell Culture. Advanced Materials 23, 1341–1345 (2011).

55. St. John, J. C. et al. The Analysis of Mitochondria and Mitochondrial DNA in Human Embryonic Stem Cells. in Human Embryonic Stem Cell Protocols vol. 331 347–374 (Humana Press, 2006).

56. Prigione, A., Fauler, B., Lurz, R., Lehrach, H. & Adjaye, J. The Senescence-Related Mitochondrial/Oxidative Stress Pathway is Repressed in Human Induced Pluripotent Stem Cells. STEM CELLS 28, 721–733 (2010).

57. Wu, J., Ocampo, A. & Belmonte, J. C. I. Cellular Metabolism and Induced Pluripotency. Cell 166, 1371–1385 (2016).

58. Berger, E. et al. Millifluidic culture improves human midbrain organoid vitality and differentiation. Lab Chip 18, 3172–3183 (2018).

59. Jiang, B. H., Semenza, G. L., Bauer, C. & Marti, H. H. Hypoxia-inducible factor 1 levels vary exponentially over a physiologically relevant range of O2 tension. American Journal of Physiology-Cell Physiology 271, C1172–C1180 (1996).

60. Huang, L. E., Gu, J., Schau, M. & Bunn, H. F. Regulation of hypoxia-inducible factor 1 is mediated by an O2-dependent degradation domain via the ubiquitin-proteasome pathway. Proceedings of the National Academy of Sciences 95, 7987–7992 (1998).

61. Greijer, A. E. The role of hypoxia inducible factor 1 (HIF-1) in hypoxia induced apoptosis. Journal of Clinical Pathology 57, 1009–1014 (2004).

62. Wang, M., Tan, J., Miao, Y., Li, M. & Zhang, Q. Role of Ca^2+^ and ion channels in the regulation of apoptosis under hypoxia. Histol Histopathol 33, 237–246 (2018).

63. Punovuori, K. et al. N-cadherin stabilises neural identity by dampening anti-neural signals. Development 146, dev183269 (2019).

64. Tietz, P. S. & Larusso, N. F. Cholangiocyte biology. Curr Opin Gastroenterol 22, 279–287 (2006).

65. Ruebner, B. H., Blankenberg, T. A., Burrows, D. A., Soohoo, W. & Lund, J. K. Development and Transformation of the Ductal Plate in the Developing Human Liver. Pediatric Pathology 10, 55–68 (1990).

66. Limaye, P. B. et al. Expression of specific hepatocyte and cholangiocyte transcription factors in human liver disease and embryonic development. Lab Invest 88, 865–872 (2008).

67. Ranga, A. et al. Neural tube morphogenesis in synthetic 3D microenvironments. Proc Natl Acad Sci USA 113, E6831–E6839 (2016).

68. Medina, J. D. et al. Functionalization of Alginate with Extracellular Matrix Peptides Enhances Viability and Function of Encapsulated Porcine Islets. Adv. Healthcare Mater. 9, 2000102 (2020).

69. Lancaster, M. A. & Knoblich, J. A. Generation of cerebral organoids from human pluripotent stem cells. Nature Protocols 9, 2329–2340 (2014).

70. Boon, R. et al. Amino acid levels determine metabolism and CYP450 function of hepatocytes and hepatoma cell lines. Nat Commun 11, 1393 (2020).

71. Roelandt, P., Vanhove, J. & Verfaillie, C. Directed Differentiation of Pluripotent Stem Cells to Functional Hepatocytes. in Pluripotent Stem Cells (eds. Lakshmipathy, U. & Vemuri, M. C.) vol. 997 141–147 (Humana Press, 2013).

72. Stuart, T. et al. Comprehensive Integration of Single-Cell Data. Cell 177, 1888–1902.e21 (2019).

73. Cao, J. et al. The single-cell transcriptional landscape of mammalian organogenesis. Nature 566, 496–502 (2019).

74. Kumar, M. et al. A fully defined matrix to support a pluripotent stem cell derived multi-cell-liver steatohepatitis and fibrosis model. Biomaterials 276, 121006 (2021).

